# Inhibition of Mycobacterium tuberculosis DosRST two-component regulatory system signaling by targeting response regulator DNA binding and sensor kinase heme

**DOI:** 10.1101/411793

**Authors:** Huiqing Zheng, Bilal Aleiwi, Edmund Ellsworth, Robert B. Abramovitch

**Affiliations:** Department of Microbiology and Molecular Genetics; Department of Pharmacology and Toxicology, Michigan State University East Lansing Michigan 48824

**Keywords:** Chemical biology, two-component regulatory systems, microbial pathogenesis

## Abstract

*Mycobacterium tuberculosis* (Mtb) possesses a two-component regulatory system, DosRST, that enables Mtb to sense host immune cues and establish a state of non-replicating persistence (NRP). NRP bacteria are tolerant to several anti-mycobacterial drugs and are thought to play a role in the long course of tuberculosis (TB) therapy. Therefore, small molecules that inhibit Mtb from establishing or maintaining NRP could reduce the reservoir of drug tolerant bacteria and function as an adjunct therapy to reduce treatment time. Previously, we reported the discovery of six novel chemical inhibitors of DosRST, named HC101A-106A, from a whole cell, reporter-based phenotypic high throughput screen. Here, we report functional and mechanism of action studies of HC104A and HC106A. RNAseq transcriptional profiling shows that the compounds downregulate genes of the DosRST regulon. Both compounds reduce hypoxia-induced triacylglycerol synthesis by ~50%. HC106A inhibits Mtb survival during hypoxia-induced NRP, however, HC104A did not inhibit survival during NRP. An electrophoretic mobility assay shows that HC104A inhibits DosR DNA binding in a dose-dependent manner, indicating that HC104A may function by directly targeting DosR. In contrast, UV-visible spectroscopy studies suggest HC106A directly targets the histidine kinase heme, via a mechanism that is distinct from the oxidation and alkylation of heme previously observed with artemisinin (HC101A). Synergistic interactions were observed when DosRST inhibitors were examined in pair-wise combinations with the strongest potentiation observed between artemisinin paired with HC102A, HC103A, or HC106A. Our data collectively show that the DosRST pathway can be inhibited by multiple distinct mechanisms.

## Introduction

*Mycobacterium tuberculosis* (Mtb) is the causative agent of tuberculosis (TB). Mtb is an intracellular pathogen that can latently infect the host without causing disease symptoms (1). During chronic infection, it can establish a dormant state known as non-replicating persistence (NRP) where Mtb modulates its metabolism in response to environmental and host immune cues, such as hypoxia, acidic pH, and nutrient starvation (2, 3). DosRST is a two-component regulatory system that regulates Mtb persistence (4-6). It consists of two sensor histidine kinases, DosS and DosT, and the cognate response regulator DosR, which regulates expression of about 50 genes in the DosRST regulon (6-8). The pathway can be induced by host intracellular stimuli, such as nitric oxide (NO), carbon monoxide (CO) and hypoxia, through DosS and DosT (9-11). DosS is an oxygen and redox sensor, whereas DosT acts an oxygen sensor (12-14). Both kinases sense ligands via the heme group, and are inactive when the heme exists as either the Met (Fe^3+^) form (DosS) or the oxy (Fe^2+^-O_2_) form (DosT) in the presence of O_2_ (13). However, hypoxic conditions activate the kinases by inducing the conversion of DosS to the ferrous form and DosT to the deoxy form. Therefore, DosS/T play overlapping and distinct roles in sensing the redox status and oxygen level of the environment to turn on the DosR pathway (11, 15).

Non-replicating bacilli are naturally tolerant to many anti-mycobacterial drugs, such as isoniazid (INH) that only kills replicating Mtb (16-18). During TB infection, the disease presents as a spectrum of replicating and non-replicating bacteria; the NRP population of bacteria are thought to be responsible, in part, for the 6-month long course of TB treatment. This long antibiotic regimen makes controlling the TB epidemic challenging and has likely contributed to the evolution of drug-resistant Mtb strains. Therefore, there exists an urgent need to identify new strategies to treat the disease, with a particular focus on discovering new ways to shorten the course of TB therapy. Targeting the DosRST pathway is a promising strategy, because *dosRST* mutants are attenuated in *in vitro* models of hypoxia-driven NRP (19) and in animal models that generate hypoxic granulomas, including non-human primates, guinea pigs, and C3HeB/FeJ mouse models of TB infection (4, 20-22). Furthermore, deletion of DosR-regulated gene *tgs1*, which is involved in triacylglycerol (TAG) synthesis, causes reduced antibiotic tolerance (23, 24). Therefore, inhibiting the DosRST pathway and killing the reservoir of NRP bacteria may function to narrow the spectrum of TB disease and shorten the course of TB therapy.

Fluorescent reporter strains can be used as tools in drug discovery to conduct targeted, whole cell phenotypic screens for pathways that play a role during pathogenesis, but have limited impact when bacteria are grown *in vitro*(25, 26). In an effort to discover the new chemical probes that inhibit Mtb persistence, we previously performed a whole cell phenotypic high throughput screening (HTS) of a >540,000 compound library using the DosRST regulon reporter strain CDC1551 (*hspX’::*GFP) (27). We discovered six compounds that inhibit the DosR-dependent, hypoxia-induced GFP fluorescence. In the previous report, we showed that the HC101, HC102 and HC103 series functioned to inhibit NRP associated physiologies, including TAG accumulation, survival during hypoxia and isoniazid tolerance. Mechanism of action studies showed that the HC101 series, composed of artemisinin and related analogs, functioned by oxidizing and alkylating the DosS and DosT heme. HC102 and HC103 did not modulate the DosS/T heme, and were instead found to inhibit sensor kinase autophosphorylation. Here, we report the characterization of two additional compounds, HC104A and HC106A. Transcriptional and biochemical analysis demonstrate that both compounds function to downregulate the DosRST pathway and inhibit persistence associated physiologies. Biochemical studies show HC104A and HC106A function by distinct mechanisms of action, with HC104A inhibiting DosR DNA binding and HC106A interacting with DosS and DosT heme to block environmental sensing. Studies examining the pair-wise interactions between the five DosRST inhibitors revealed synergistic interactions, including strong potentiating interactions of artermisinin and HC106A. Structure activity relationship studies of HC106 identified functional groups of HC106A that are required for activity and enabled optimization of HC106 potency to nanomolar effective concentrations against whole cell Mtb.

## Results

### Inhibition of the DosR regulon by HC104A and HC106A

Characterization studies were undertaken with two putative DosRST regulon inhibitors, HC104A and HC106A (Fig. 1a) (27). Half-maximal effective concentration (EC_50_) studies using the CDC1551 (*hspX’::GFP*) DosRST-dependent fluorescent reporter strain, show that HC104A and HC106A inhibit DosRST-dependent GFP florescence with EC_50_ values of 9.8 μM and 2.48μM, respectively (Fig. 1b and 1c).

**Figure 1.**
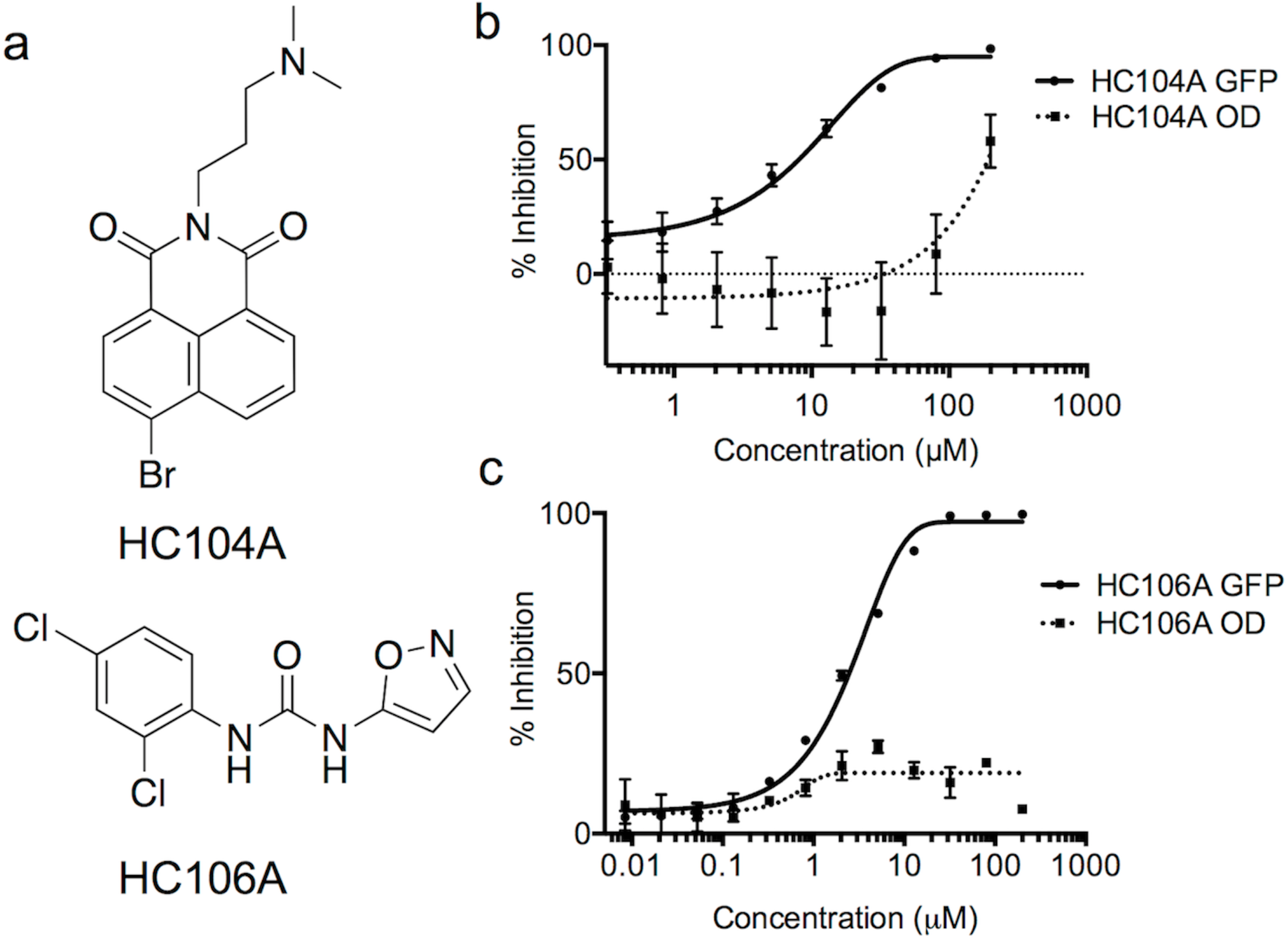
HC104A and HC106A inhibit DosRST reporter fluorescence. (**a**) Chemical structures of HC104A and HC106A. HC104A (**b**) and HC106A (**c**) inhibited DosR-driven GFP fluorescence signal in a dose-dependent manner, while having minimal impact of Mtb growth. The EC_50_ values of fluorescence inhibition for HC104A and HC106A are 9.8 μM and 2.5 μM, respectively.

The compounds have minimal impact on Mtb growth, suggesting they are also potential DosRST inhibitors, as DosRST is not required for survival under the conditions of mild hypoxia used in the reporter-based assay. RNA-seq-based transcriptional profiling was undertaken to determine if the DosRST regulon was inhibited by the compounds. Mtb was treated with 40 μM HC104A, HC106A or dimethyl sulfoxide (DMSO) control for 6 d in a standing flask, and following incubation RNA was extracted, sequenced and analyzed for differential gene expression relative to the DMSO control. As a control for the DosR regulon, transcriptional profiling was also previously conducted on a DMSO treated CDC1551(Δ*dosR*) mutant strain (27). The transcriptional profiles showed that the genes strongly repressed by HC104A and HC106A (>2-fold; q<0.05) are from the *dosR* regulon (Fig. 2a-c, Supplementary Dataset 1, 3).

**Figure 2.**
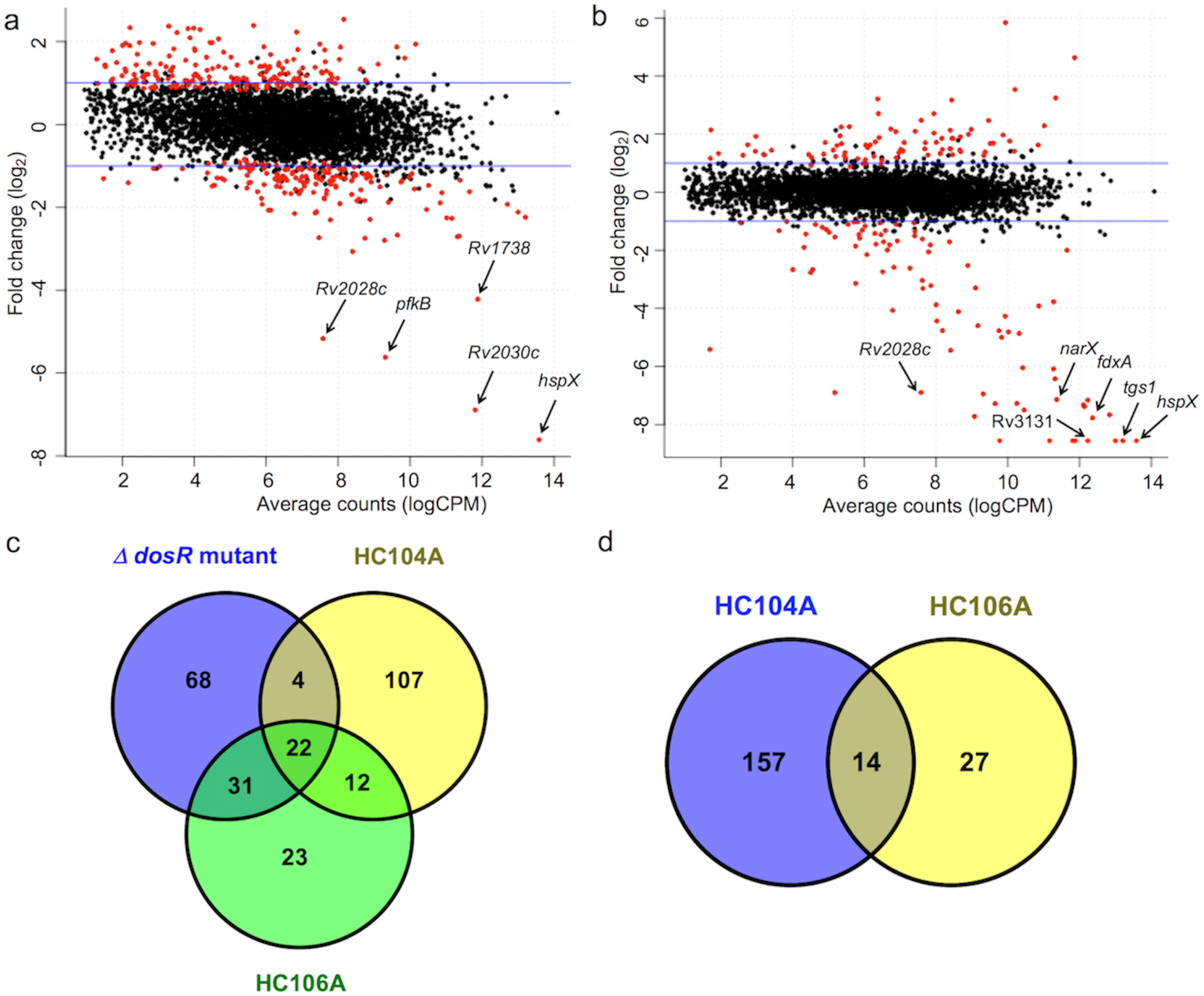
Transcriptional profiling shows HC104A and HC106A inhibited DosR regulon during hypoxia. Magnitude-amplitude plots showing differential gene expression of Mtb cells treated with 40 μM HC104A (**a**) or HC106A (**b**). The labeled genes represent selected genes that belong to the DosRST regulon. The red dots represent genes with significant differential expression, q<0.05. **(c)** A Venn diagram for the downregulated genes (>2-fold; q*<* 0.05) of WT CDC1551 treated with HC104A or HC106A compared to that of CDC1551 (Δ*dosR*). (**d**) Venn diagram for downregulated genes (>2-fold; q< 0.05) of CDC1551 (Δ*dosR*) treated with HC104A or HC106A.

HC106A exhibited a remarkably strong reduction of gene expression, with transcripts for *tgs1* and *hspX* being almost undetectable by RNA-seq following HC106A treatment. Interestingly, while HC106A broadly inhibited genes of the DosRST regulon, HC104A only strongly inhibited part of the DosR regulon, with the strongest inhibition reserved for *hspX*, the promoter used to drive reporter fluorescence in the screen. These RNA-seq results were validated by semi-quantitative RT-PCR, with HC104A causing downregulation of *dosR, hspX*, and *tgs1 in vitro* by 6-, 570-and 13-fold, respectively; whereas HC106A downregulated these three genes by 49-, 1360-, and 1424-fold, respectively (Supplementary Fig. 1), with *hspX* and *tgs1* transcripts being below the level of detection by qRT-PCR.

Comparisons of transcriptional profiles from the inhibitor treated wild type (WT) Mtb strain to a CDC1551(Δ*dosR*) mutant strain showed that there are a total of 26 genes and 53 genes from *dosR* regulon inhibited by HC104A and HC106A, respectively. Notably, HC104A and HC106A caused an additional 119 genes and 35 genes to be repressed that were not repressed in the CDC1551(Δ*dosR*) mutant strain (Fig. 2c). This observation suggested that these two compounds exhibit some DosR-independent activities. To confirm the specificity of the compounds, RNA-seq was also performed on CDC1551(Δ*dosR*) mutant background (Supplementary Dataset 2-3) treated with HC104A or HC106A. This analysis identified 171 genes and 51 genes that are downregulated (>2-fold; q<0.05) by HC104A and HC106A, respectively (Fig. 2d). This finding indicates that HC104A and HC106A impact other targets beside DosR regulon, with HC106A showing greater on-target specificity than HC104A. Based on these findings, we conclude that: 1) HC106A strongly and specifically inhibits the DosRST regulon; and 2) HC104A strongly inhibits a portion of the DosRST regulon, with several notable off-target activities.

To assess the impact of the inhibitors on the DosRST pathway on intracellular Mtb, murine bone marrow-derived mouse macrophages were infected with Mtb and treated with 40 μM HC104A and HC106A for 48 h. Total bacterial RNA was isolated and analyzed by RT-PCR for *hspX* and *tgs1* gene differential expression. The results demonstrate that the induction of *hspX* and *tgs1* were inhibited 185-and 10-fold by HC104A and 6-and 4-fold by HC106A, respectively (Fig. 3a). These finding confirm that HC104A and HC106A can access Mtb inside the macrophage, however, the reduced repression of the pathway by HC106A as compared to broth culture, suggests that the molecule may not be able to efficiently target intracellular Mtb.

**Figure 3.**
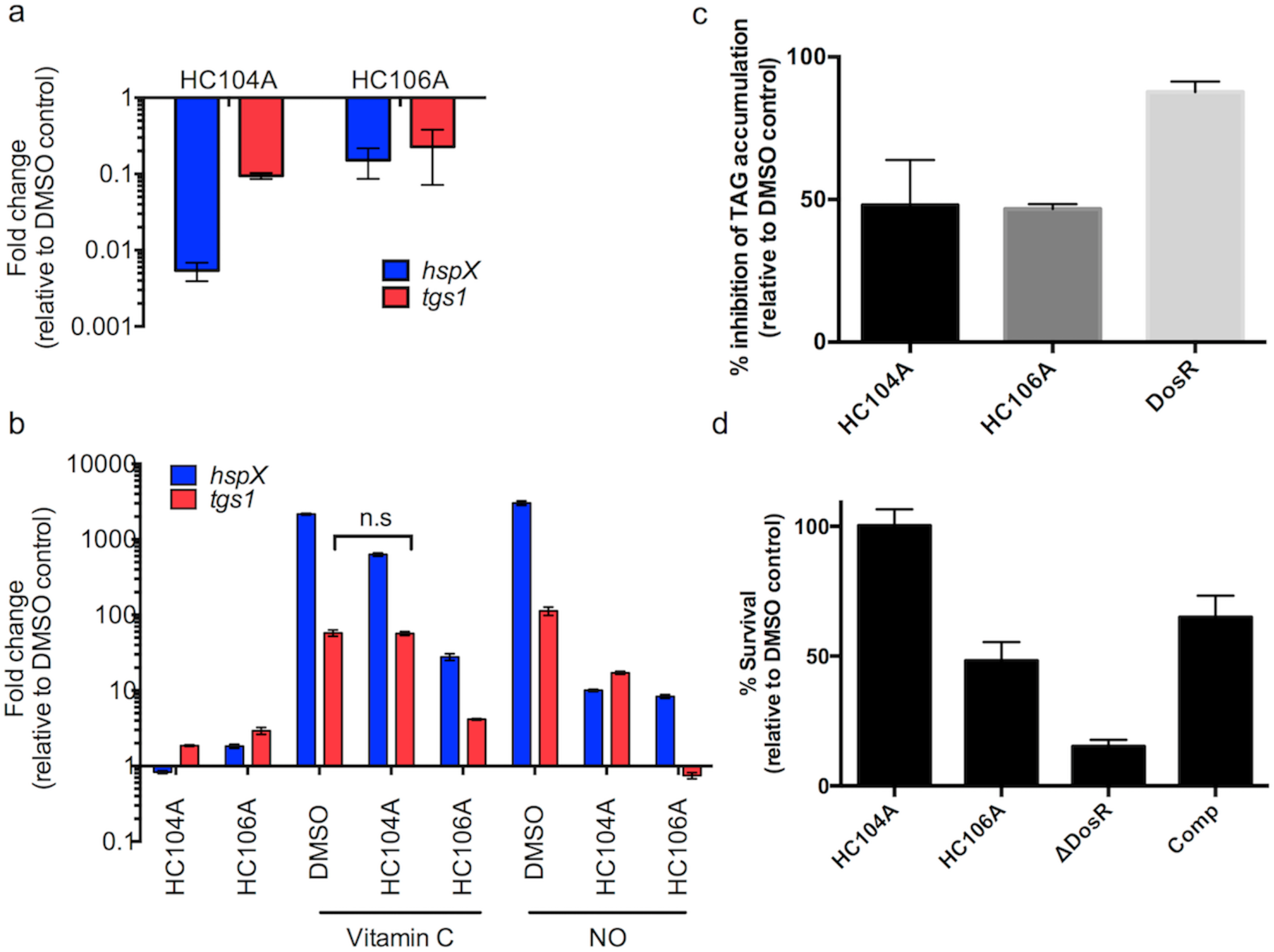
Inhibition of DosR regulon and persistence-associated physiologies by HC104A and HC106A. (**a**) Inhibition of DosR regulon in murine macrophages infected with Mtb and treated with HC104A and HC106A for 48 h. Bacterial RNA was isolated after incubation, and the differential gene expression of *hspX* and *tgs1* bone-marrow derived were quantified by qRT-PCR. The error bars represent the standard derivation of three biological replicates. (**b**) HC104A and HC106A inhibit DosR regulon induction by vitamin C and NO. Cells were pretreated with the compounds or DMSO for 24 h, and induced with vitamin C or NO for 2 h. Total bacterial RNA was isolated, and the transcripts of DosR-regulated genes, *hspX* and *tgs1*, were quantified by qRT-PCR. The difference in the drug treated samples compared to DMSO treated samples in response to vitamin C or NO are significant with a p-value <0.001 based on a t-test, except the one marked as non-significant (n.s.). The error bars represent the standard deviation of three replicates. The experiment was repeated twice with similar results. (**c**) Inhibition of TAG accumulation of Mtb treated with HC104A or HC106A. Mtb cells were treated with 40 μM of the compounds and labeled with [1,2-^14^C] sodium acetate for 6 d. Total lipid was isolated and analyzed by TLC. TAG accumulation was quantified from the TLC. The error bars represent the standard derivation of two biological replicates. (**d**) Mtb cell survival during NRP when treated with HC104A or HC106A during NRP. Mtb cells were pretreated with 40 μM of compounds for 48 h in an anaerobic chamber, and continued incubation for 10 d. Surviving bacteria were enumerated on 7H10 agar. The error bars represent the standard derivation of three biological replicates. The experiment was repeated twice with similar results.

The DosRST pathway is also induced by redox signals such as vitamin C and NO (27). To examine whether HC104A and HC106A can repress the induction of DosRST pathway by vitamin C or NO, Mtb cells were pretreated with HC104A or HC106A for 24 h followed by vitamin C or NO induction for 2 h. The expression of DosR-regulated genes (*hspX* and *tgs1*) was examined by real time-PCR. Vitamin C and DETA-NONOate (NO donor) strongly induced *hspX* and *tgs1* as previously reported ((27), Fig. 3b). For instance, vitamin C induced *hspX* and *tgs1* by 2162-and 58-fold, respectively; whereas DETA-NONOate upregulated *hspX* and *tgs1* 3024-and 113-fold, respectively (Fig. 3b). Mtb cells pretreated with HC106A showed strong inhibition of *hspX* and *tgs1* induction by vitamin C and DETA-NONOate. For example, HC106A inhibited the *hspX* and *tgs1* transcripts by 78-and 14-fold following vitamin C treatment, respectively, and 362-and 151-fold following DETA-NONOate treatment. Following vitamin C treatment, HC104 showed inhibition of *hspX* by 3.4-fold and no effect on *tgs1*, or 302-and 6.6-fold inhibition of *hspX* and *tgs1* following DETA-NONOate treatment. These findings show that HC104A and HC106A act as inhibitors of the DosRST pathway in response to redox signals in addition to hypoxia.

### Inhibition of Mtb persistence-associated physiologies

Mtb accumulates TAG during hypoxia (27-29). DosR directly regulates *tgs1*, which encodes a TAG synthase that is involved in the last step of TAG biosynthesis and is required for TAG accumulation during hypoxia (30). Transcriptional profiling showed that HC104A and HC106A repress expression of *tgs1*. Based on this transcriptional profiling, we hypothesized these two compounds may inhibit TAG biosynthesis during NRP. To test our hypothesis, Mtb cells were radiolabeled with ^14^C-acetate and treated with HC104A or HC106A for 6 d. Lipids were isolated and analyzed by thin layer chromatography (TLC). As previously observed, DMSO treated CDC1551(Δ*dosR*) mutant displayed a strong (87%) reduction of TAG accumulation as compared to DMSO treated WT (Fig. 3c and Supplementary Fig. 2). Mtb cells treated with HC104A or HC106A showed a ∼50% reduction of TAG accumulation, supporting our hypothesis that the compounds can inhibit TAG biosynthesis.

DosRST has been previously reported to be required for survival during NRP, where deletion of DosR causes greatly reduced survival during prolonged hypoxic stress (19). The impact of HC104A and HC106A on Mtb survival during NRP was examined using the hypoxic shift-down model (27). Mtb survival was examined following 10 d of treatment with the compounds at 40 μM. The Δ*dosR* mutant control had 15% survival relative to DMSO, and was partially complemented, supporting the proposal that survival during hypoxia is DosR dependent. Mtb cells treated with HC106A displayed 50% survival relative to DMSO control (Fig. 3d), whereas, HC104A had no impact on Mtb survival during NRP, an observation that suggests the portion of the DosR regulon inhibited by HC104A is not essential for survival during NRP.

### HC104A inhibits DosR DNA binding

There are several potential targets of HC104A and HC106A to directly inhibit DosRST signaling, including: 1) direct inhibition of DosS/T sensor kinase activity, 2) modulation of the heme in the sensor, or 3) inhibition of DosR binding of DNA. To investigate the biochemical mechanisms of action of HC104A and HC106A, inhibition of DosS autophosphorylation was initially evaluated. The DosS protein was treated with different concentrations of HC104A and HC106A from 10 μM to 40 μM, or with 40 μM HC103A as a positive control that was previously discovered to be a DosS/T autophosphorylation inhibitor (27). As previously observed, HC103A strongly inhibited DosS autophosphorylation, but HC104A and HC106A had no inhibitory activity (Supplementary Fig. 3). This suggests HC104A and HC106A are not directly inhibiting DosS/T autophosphorylation activity.

Next, a UV-visible spectroscopy assay was employed to investigate if HC104A targets to the heme of the sensor kinase DosS. Treatment of DosS protein with the reducing agent dithionite (DTN) caused a shift of the Soret peak from 403 nm to 430 nm as previously described (13, 27). Addition of HC104A to reduced DosS did not shift the peak to the oxidized position, suggesting that HC104A does no modulate DosS heme redox (Supplementary Fig. 4). Together, these data indicate that HC104A does not inhibit DosRST signaling by targeting sensor kinase autophosphoryation or heme and supported examining DosR as a potential target.

**Figure 4.**
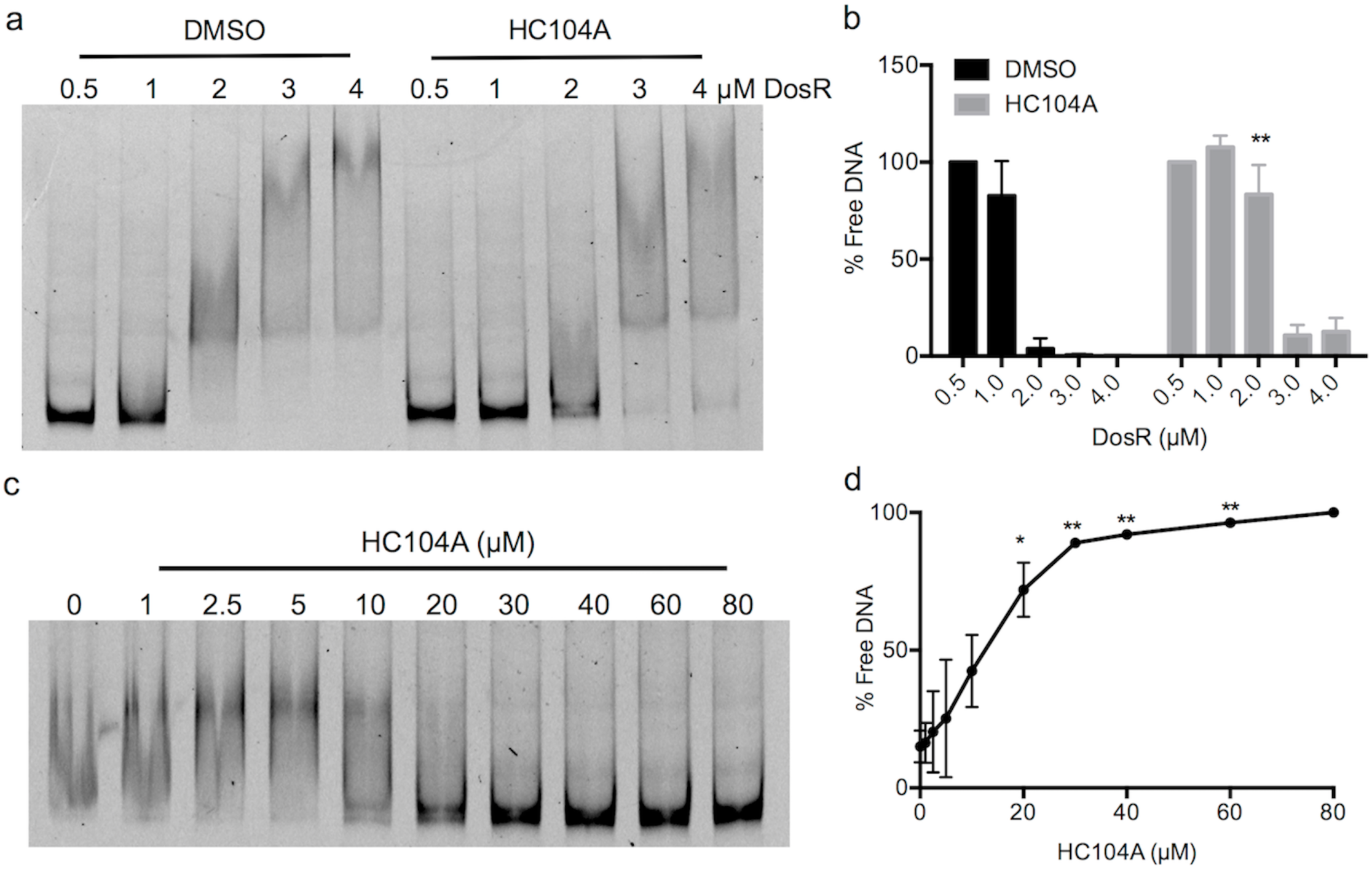
Inhibition of DosR DNA-binding by HC104A. (**a**) DosR protein ranging from 0.5 μM to 4 μM was treated with DMSO or 40 μM HC104A and binding to the *hspX* promoter was examined by EMSA. HC104A inhibits DosR DNA binding at 2 μM concentration. (**b**) The free DNA of each reaction was quantified in ImageJ and the percentage of free DNA was normalized using reactions containing 0.5 μM DosR as 100% free DNA. Differences between reactions containing 2 μM DosR treated with DMSO or HC104A are significant (\*\**P* value <0.005 based on a *t*-test). The error bars represent the standard derivation of two biological replicates. (**c**) Dose-dependent impact of HC104A on DosR DNA binding. DosR protein at 2 μM was treated with HC104A at concentrations from 1 μM to 80 μM. (**c**) The free DNA of each reaction was also quantified in ImageJ with the percentage of free DNA is normalized using the reaction containing 80 μM HC104A as 100% free DNA. The differences between treated reactions as compared to DMSO control are significant. (\**P* value <0.05 and \*\**P* value <0.005 based on a *t*-test). The error bars represent the standard derivation of two biological reps.

Inspection of the HC104A structure revealed it had significant similarity to the compound virstatin (31). Virstatin is an anti-virulence compound that inhibits *Vibrio cholera* cholera toxin production by targeting the transcription regulator ToxT (31). Virstatin inhibits ToxT protein dimerization and subsequently interferes with DNA binding, thereby inhibiting the transcription of downstream genes involved in toxin production. Based on the similarity of chemical structure between HC104A and virstatin, we hypothesized that HC104A may be targeting DosR, and interfering with DNA binding. To test this hypothesis, an electrophoretic mobility shift assay (EMSA) was employed to investigate the impact of HC104A on DosR DNA binding. Recombinant DosR protein, ranging from 0.5 μM to 4 μM, was treated with 40 μM HC104A or a DMSO control and tested for binding to fluorescently labeled *hspX* promoter DNA. In the DMSO treated control, DosR bound promoter DNA beginning at a concentration 2 μM DosR protein (Fig. 4a). Treating the reaction containing 2 μM DosR protein with HC104A significantly inhibited DNA binding by ∼22-fold compared to DMSO control (Fig. 4b). To further characterize the impact of HC104A on DosR binding of DNA, a dose-response study was performed. Reactions containing 2 μM recombinant DosR proteins were treated with different concentrations of HC104A or virstatin ranging from 1 – 80 μM. HC104A inhibited DosR binding of DNA beginning at 10 μM HC104A. The fraction of free DNA increased as HC104A concentration increased (Fig. 3.4c). For example, the fraction of free DNA was 72%, 89%, 92%, 96% and 100% for 20 μM, 30 μM, 40 μM, 60 μM and 80 μM HC104A, respectively, whereas DMSO control had 15% free DNA (Fig. 4d). Thus, HC104A significantly inhibits DosR-DNA binding in a dose-dependent manner. Reactions treated with virstatin had no impact on DosR binding of DNA (Supplementary Fig. 5a). Consistent with these observations, virstatin did not have any impact on DosRST signaling in the whole cell Mtb fluorescence reporter assay (Supplementary Fig. 5b). These findings support the hypothesis that HC104A may function by inhibiting DosR DNA binding activity and has an activity that is distinct from the related molecule virstatin.

**Figure 5.**
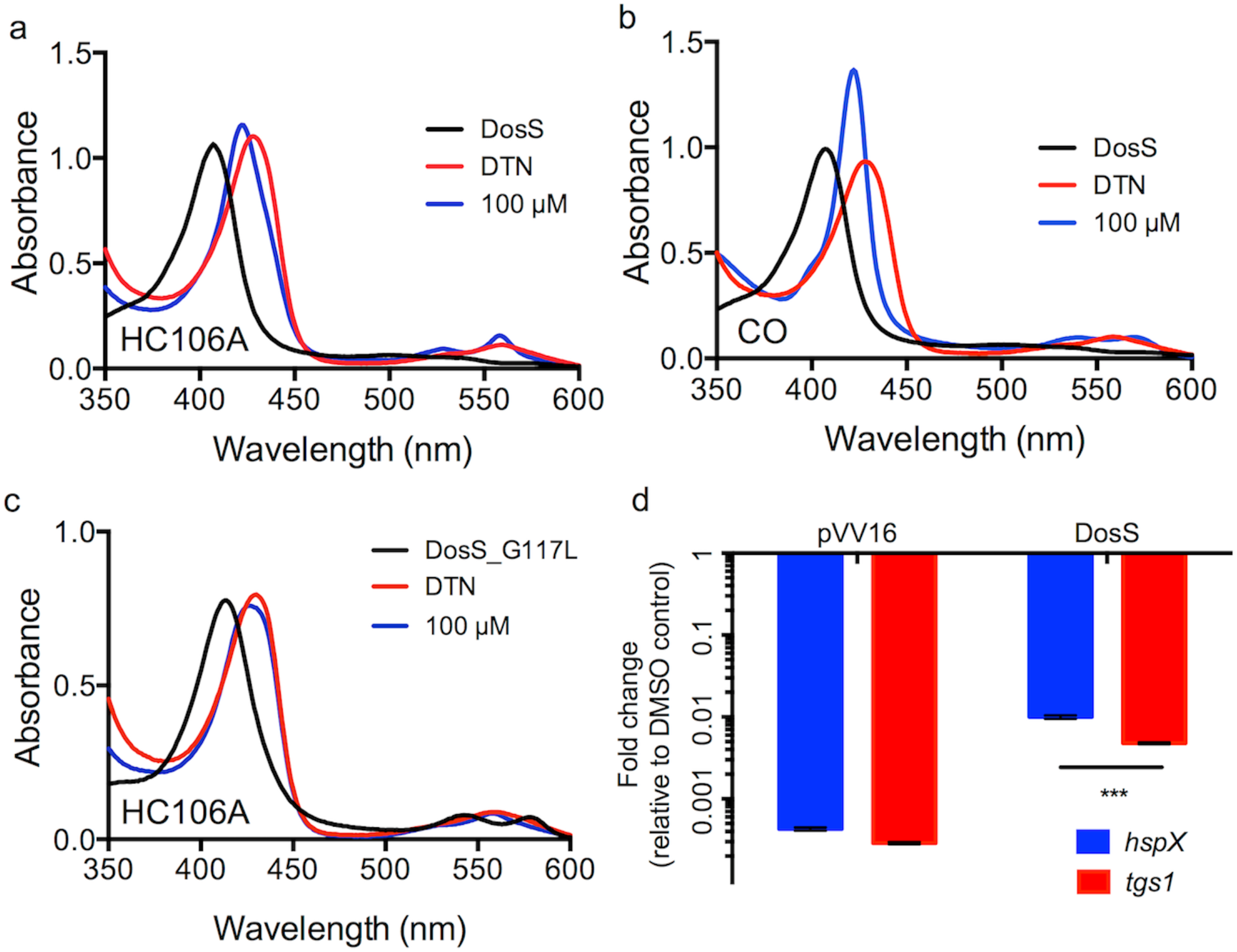
Interactions between HC106A and DosS heme. WT DosS protein was treated with dithionite (DTN) and then 100 μM HC106A (**a**) or 100 μM pf CORM-2 (a CO donor) (**b**). The UV-visible spectra of the two treatments exhibited a shift of the Soret peak to a common position of 422 nm. (**c**) DosS with a G117L amino acid substitution, that is predicted to block the heme exposing channel, provides resistance to HC106A. The spectrum of the mutant protein did not change, after HC106A treatment, indicating resistance to HC106A. (**d**) Overexpression of DosS protein promotes resistance to HC106A treatment in Mtb. Mtb cells with the pVV16 empty vector or the DosS overexpression plasmid were treated with 20 μM HC106A for 6 d. Bacterial RNA was isolated for analysis of the differential gene expression of *hspX* and *tgs1* and analyzed by qRT-PCR. Overexpression of DosS caused 23-and 16.5-fold increase of *hspX* and *tgs1* transcripts, respectively, compared to the empty vector control (\*\*\**P* value <0.0001 based on a *t*-test). The error bar represents the standard derivation of the mean for three technical replicates. The experiments were repeated twice with a similar result.

### HC106A modulates DosS heme

DosS and DosT have a channel that exposes the heme to the environment and enables interactions with gases (32, 33). This channel is an Archilles heel that can be targeted by small molecules. Previously, it was shown that the artemisinin modulates DosS/T by oxidizing and alkylating heme carried by the kinases (27). UV-visible spectroscopy studies were conducted to examine if HC106A modulated DosS heme. Recombinant DosS was purified from *E. coli*, degassed and the change of DosS heme spectrum was monitored under anaerobic conditions by UV-visible spectroscopy. Treating DosS with the reducing agent dithionite (DTN) caused the Soret peak to shift to 430 nm as shown previously (13, 27). HC106A was added to the reaction following DTN treatment to observe the impact on the DosS heme UV-visible spectrum. HC106A caused the DosS Soret peak to immediately shift to 422 nm, where the peak was stably maintained for 2 h (Fig. 5a). This spectrum shift is different from artemisinin, where under identical conditions, artemisinin causes the DosS Soret peak to gradually shift back to the oxidized state at 403 nm (27). These findings show that HC106A may also interact with sensor kinase heme, but via a mechanism that is distinct from artemisinin-heme interactions.

The Soret peak at 422 nm is consistent with previously described spectra that are observed when DosS heme interacts with NO or CO (13). To confirm this observation, DosS was treated with 100 μM CORM-2 (a CO donor) which caused a shift of the Soret peak to 422 nm, similar to what was observed for HC106A (Fig. 5b). This finding supports a hypothesis that HC106A may also be directly binding to the heme. Notably, CO activates DosS kinase function, whereas HC106A functions to inactivate the regulon, demonstrating that the impact of heme binding by CO or HC106A has differing impacts on the sensor kinase switch.

Amino acid substitutions in the channel exposing the DosS heme to the environment, such as DosS E87L or G117L, can limit access of artemisinin to modulate heme (27). To support the hypothesis that HC106A directly targets DosS heme, we tested the impact of these amino acid substitutions on HC106A/DosS heme interactions. Treating DosS(E87L) with HC106A exhibited a profile similar to wild type DosS with the Soret peak shifting to 422 nm (Supplementary Fig. 6). However, DosS(G117L) had no change to the overall spectrum after HC106A treatment (Fig. 3.5c). This finding indicates that DosS(G117L) is resistant to HC106A, and confirms that HC106A accesses the heme via a similar mechanism as artemisinin.

**Figure 6.**
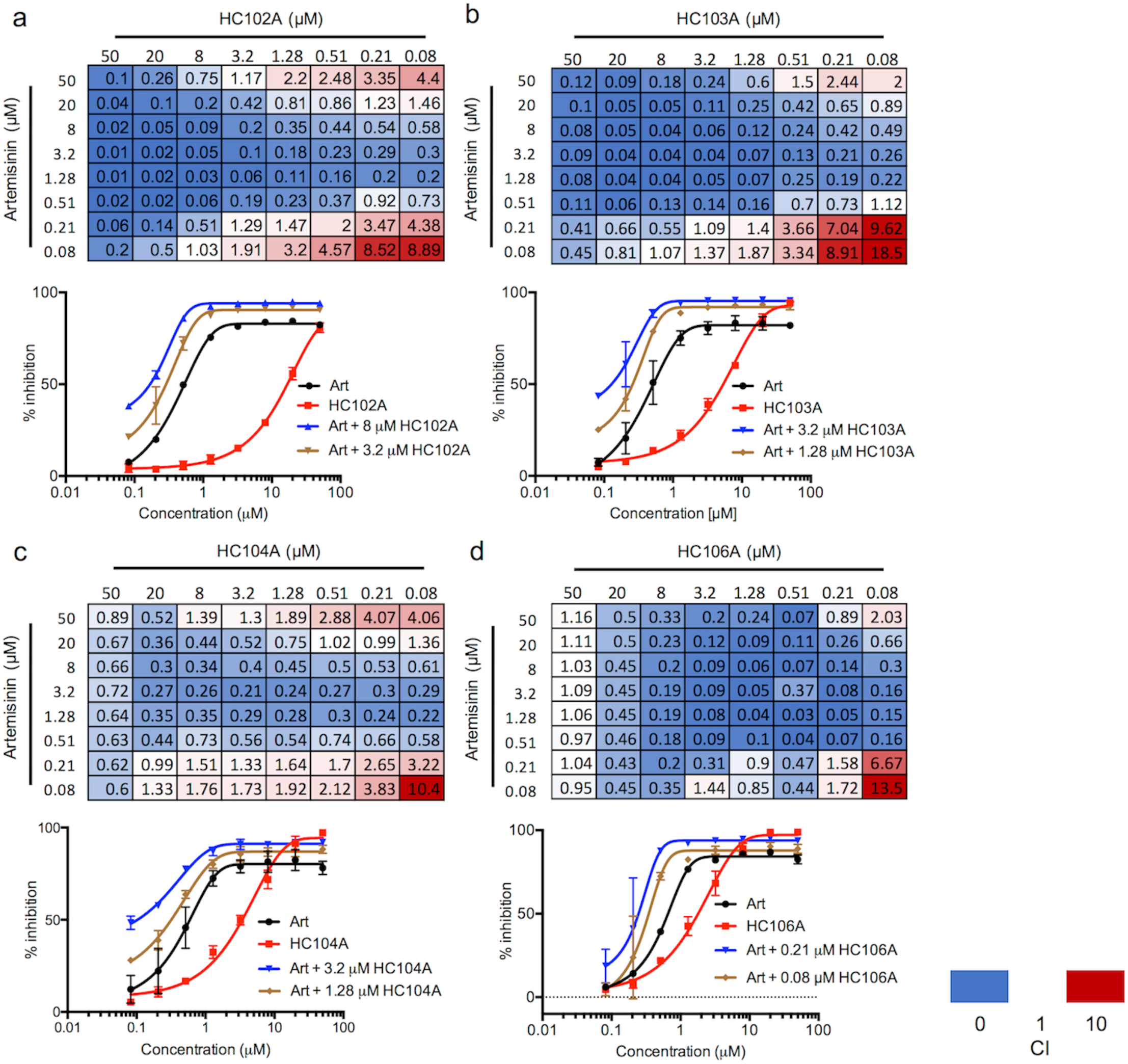
Synergistic interactions between DosRST inhibitors. CDC1551 (*hspX’::GFP*) was treated with pairwise combinations of two compounds at concentrations of 50 μM to 0.08 μM. GFP fluorescence was measured and used to calculate percentage inhibition. The data were analyzed in the CompuSyn software to determine the combination index (CI) for the panel of each drug combination, including (**a**) artemisinin and HC102A; (**b**) artemisinin and HC103A; (**c**) artemisinin and HC104A; and (**d**) artemisinin and HC106A. Example EC_50_ curves are presented with individual compounds or a selected synergistic combination to illustrate the potentiating interactions.

To confirm DosS is a target of HC106A in Mtb, we examined the impact of overexpressing DosS protein in Mtb. If DosS is the target of HC106A, overexpression may reduce the effectiveness of HC106A. WT DosS protein was constitutively expressed from the *hsp60* promoter in Mtb. The vector control showed that both *hspX* and *tgs1* genes were downregulated by HC106A by 2331-and 3470-fold, respectively (Fig. 5d). Overexpressing DosS provided significant resistance to HC106A, with *hspX* and *tgs1* showing 23-and 16.5-fold less inhibition, respectively, relative to the empty vector control This observation of resistance in Mtb is consistent with the biochemical data supporting the view that DosS is a direct target of HC106A.

### Synergistic interactions between DosRST inhibitors

With multiple distinct inhibitory activities of the HC101A, HC102A, HC103A, HC104A and HC106A, we sought to examine if potentiating or antagonistic interactions existed between the molecules, when targeting the DosRST pathway. To examine these interactions, checkboard assays were performed with pairwise comparisons of artemisinin (HC101A), HC102A, HC103A, HC104A and HC106A. CDC1551 (*hspX’::*GFP) was treated with combinations of two compounds ranging from 50 μM to 0.08 μM in 96-well plates. DosR-driven GFP fluorescence and optical density were measured following 6 d of treatment. The Combination Index (CI) was calculated for each drug pair based on the Chou-Talalay method in the CompuSyn software package (34, 35), where CI values of < 1, = 1 and >1 indicate synergistic, additive or antagonistic interactions, respectively. Among all 64 compound pairs, artemisinin combined with HC102A, HC103A, HC104A and HC106, showed 46, 49, 41, and 50 combinations that have CI <1, respectively (Fig. 3.6). Notably, some CI values are below 0.1 when artemisinin was paired with HC102A, HC103A or HC106A combinations. Example dose response curves illustrate these synergistic interactions (Fig. 6). Several other pairwise comparisons also demonstrated synergy (Supplementary Fig. 7), however, in general, these interactions had CI between 1 and 0.1, supporting weaker synergistic interactions, as compared to combinations with artemisinin. Overall, these studies provide the evidence that the inhibitors function by distinct mechanisms and may be combined to improve potency.

### Structure-activity relationship study for HC104A and HC106

We conducted a catalog search for HC104 and HC106 analogs and obtained 10 commercial analogs for each series to define initial structure activity relationships (SAR). For HC104A we observed that a bromine in the 5-position is required for activity and that the R2 dimethylamine group is not required (Supplementary Table 1). For example, HC104B is identical to HC104A except for the removal of the bromine (R1), which results in a complete loss of activity in the whole cell assay. Whereas, replacement of the R2 group with a methyl (HC104G) results in an active compound, although ∼5-fold less active than HC104A. Although not highly potent, its ligand efficiency, cLogD and druglikeness are in the range of what would be considered acceptable to good as a starting point for further manipulation. For HC106A (Table 1), catalog SAR work led to new understandings of the nature of the series. We first found that the simple removal of an ortho chloro on the “A” ring of HC106A leads to ∼ 2-fold enhanced activity, with an EC_50_ in the whole cell Mtb assay for DosRST inhibition of 1.33 µM (HC106F). It was also found that the use of an alternative isomer of the isoxazole had no detectable activity (HC106C).

**Table 1.**
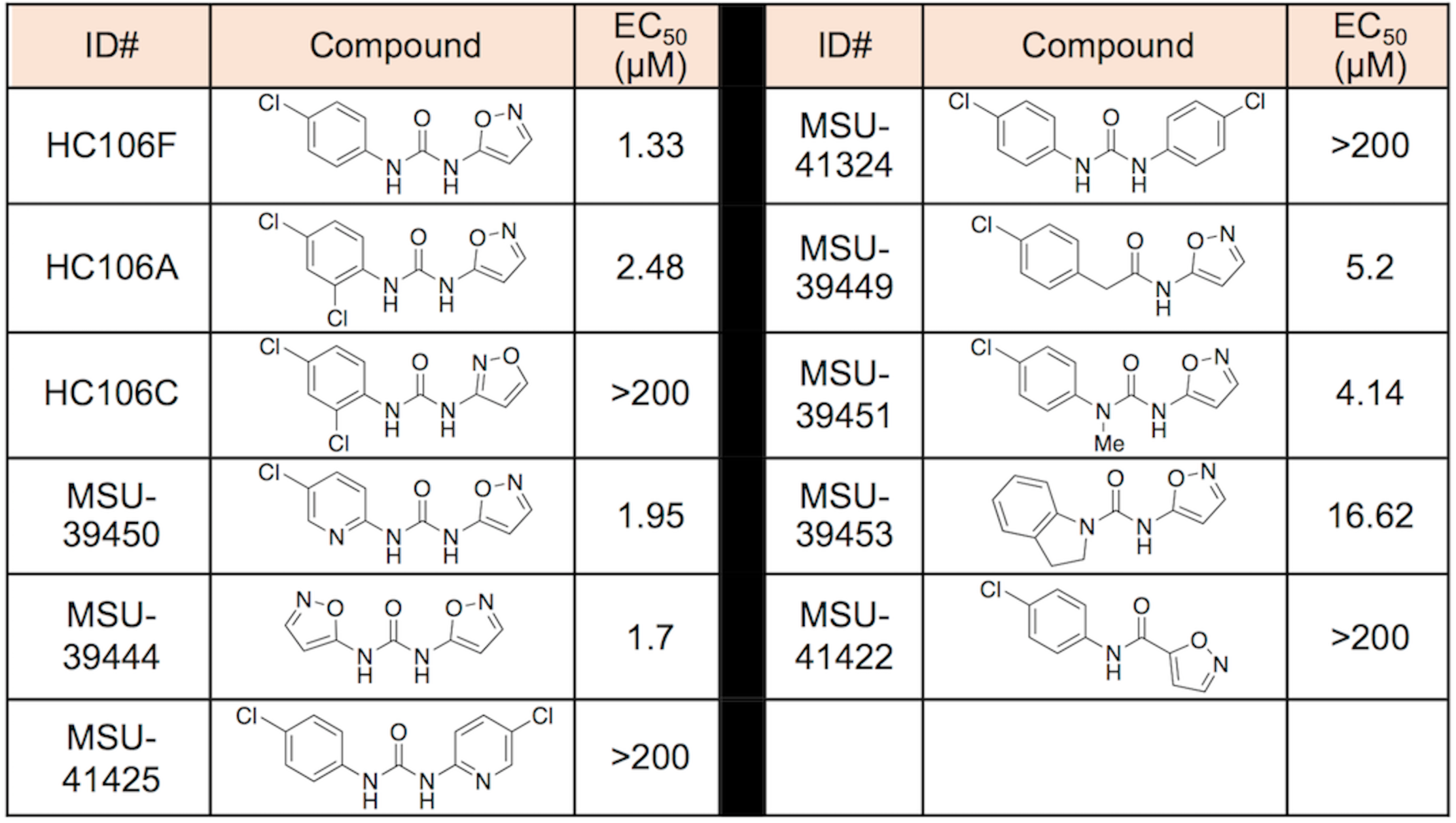
Initial SAR studies of the HC106 series. The HC106 analogs with different R-groups were synthesized or purchased. The reporter strain CDC1551 (*hspX’::GFP*) was tr6 mtreated with across doses of each analog from 200 μM to 0.328 μM. The EC_50_ values of fluorescence inhibition calculated for each analog to determine their potency.

| ID# | Compound | EC<sub>50</sub><br>(μM) | ID# | Compound | EC<sub>50</sub><br>(μM) |
| --- | --- | --- | --- | --- | --- |
| HC106F |  | 1.33 | MSU-41324 |  | >200 |
| HC106A |  | 2.48 | MSU-39449 |  | 5.2 |
| HC106C |  | >200 | MSU-39451 |  | 4.14 |
| MSU-39450 |  | 1.95 | MSU-39453 |  | 16.62 |
| MSU-39444 |  | 1.7 | MSU-41422 |  | >200 |
| MSU-41425 |  | >200 |  |  |  |

To further understand the SAR of the HC106 series, additional analogs were synthesized to examine the need of the central urea functionality and whether modifications can be tolerated (Table 1). A pyridyl analog (MSU-41425), designed to replace the isoxazole also demonstrated no activity as was the symmetrical 4-chloroaniline derived urea (MSU-41324). However, the bis-isoxazole urea (MSU-39444) provides an EC_50_ of 1.7 μM, indicating that the isoxazole is important for function. Isoxazoles are unique among heterocycles in that they exist in multiple tautomeric forms as supported by initial NMR studies (36). We next explored the need of one of the-NHs of the urea, capping it with a methyl (MSU-39451), integrating it into a ring for conformational restriction (MSU-39453), and replacing with a methylene unit (MSU-39449). In all cases, reduced activity (0.5 - 1 log) was observed but not all activity was lost. Thus, HC106A is a potent whole cell inhibitor of the DosRST pathway, with flexibility to be improved via SAR.

To further test the SAR, we conducted a Topliss Tree evaluation of the “A-ring” aniline (Table 20)(37). To reliably prepare the derivatives, we explored and established a general preparation (Supplementary Fig. 8). This route is preferred relative to alternative approaches for its cleanliness, yields and ease of purification, usually by trituration. It is also anticipated that it will allow access to future derivatives. Using HC106F and HC106A as starting points, we prepared the 3,4-diclorochloro and 3-chloro derivatives (MSU-39452 and MSU-39445, respectively). Both the 3-and 4-chloro derivatives demonstrated greater activity than 3,4-dichloro (MSU-39452). We found that replacing the 4-chlorophenyl ring with pyridyl analogs (MSU-39448 and MSU-39450) lead to similar activity. Focusing on 4-position derivatives, we found that fluoro (MSU-39446), bromo (MSU-41464) and methoxy (MSU-39447), as electron p-orbital donating substituents, also lead towards incrementally increased activity. Para-t-butyl phenyl (MSU-41442), provided slightly diminished activity. Electron withdrawing substituents, such as 4-CO_2_Me (MSU-4165), 4-trifluoromethyl (MSU-41463) and biphenyl (MSU-41443) saw activity similar to the 4-chlorophenyl derivative (HC106F). Overall, several analogs were discovered with significantly ∼4-fold enhanced potency, with several inhibitors having whole cell DosRST inhibitory EC_50_ below 1 µM, including EC_50_ of 0.63 µM, 0.61 µM, 0.54 µM and 0.75 µM for MSU-33189, -39447, -39446, and -39455, respectively. The parent analog HC106A had an EC_50_ of 2.48 µM.

There appears to be little sensitivity, positive or negative, for electron-drawing substituents other than the biphenyl derivative (MSU-41443), which is likely the result of negative steric interactions in the binding domain. Additional derivatives (replacement of dichlorophenyl) were also prepared to further probe the size and nature of the binding domain. We began by replacing the chlorophenyl ring of HC106F with benzyl-(MSU-41462), isobutyl-(MSU-41542), cyclopentyl-(MSU-41546) and cyclohexyl-(MSU-42002) analogs. All analogs demonstrated similar activities, relative to HC106F and the simple phenyl analog (MSU-33189), suggesting flexibility of fragments that could bind in this domain.

Kinetic solubility assays were conducted for selected analogs and all exhibited excellent aqueous solubility greater than >100 µM, except for MSU-41443 (Table 2). This finding shows that the urea group present in the HC106A does not have a detrimental impact on HC106 aqueous solubility. All of the tested deriviatives also demonstrated favorable mouse microsomal stability, including. Overall, the nanomolar whole cell potency, flexible SAR, good microsomal stability and excellent solubility, confirm that the HC106 series is a suitable series for continued optimization to identify a drug-like lead.

**Table 2.**
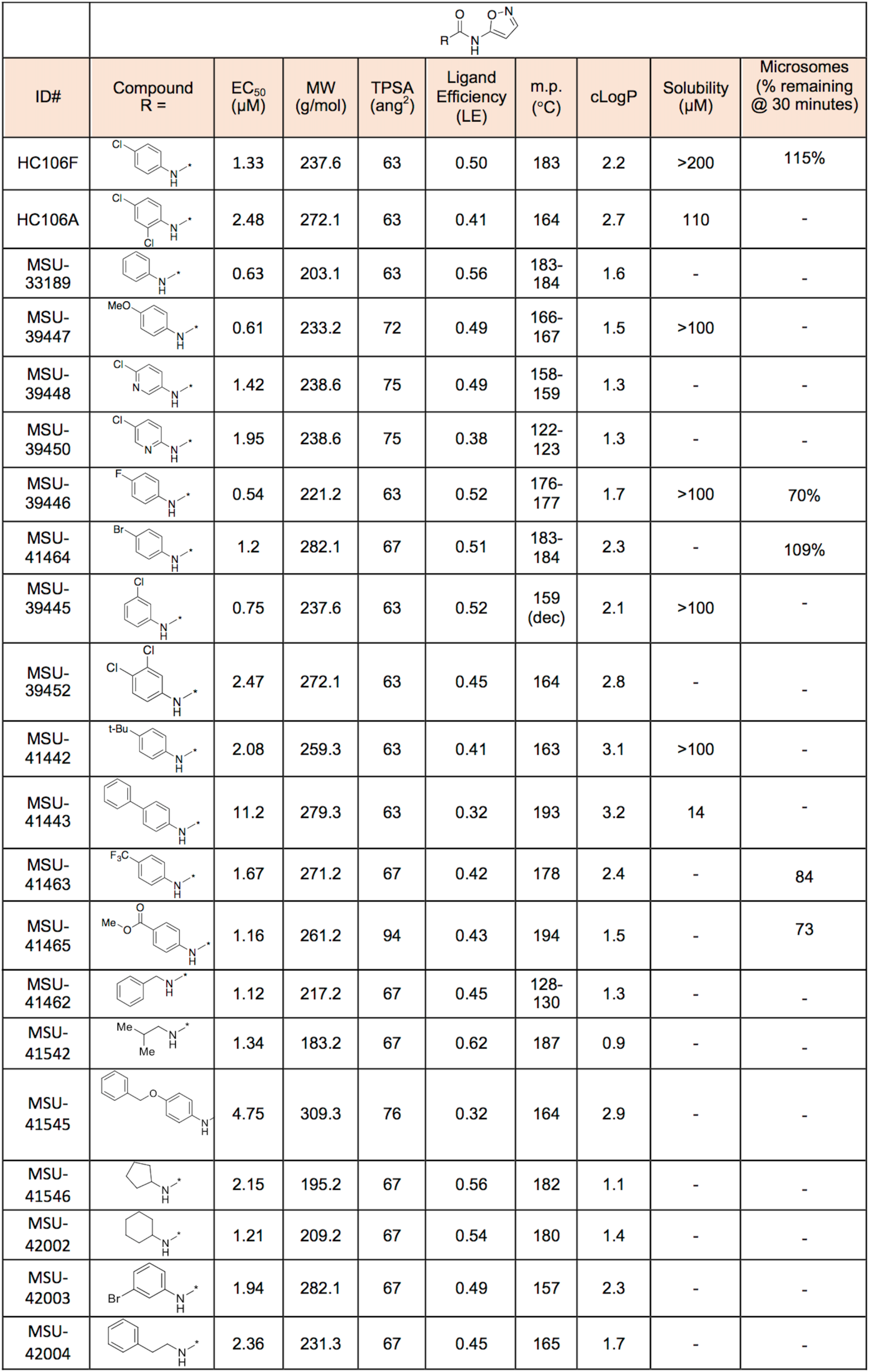
Early SAR studies of “A-ring” analogs of HC106. The HC106 analogs with different R-groups were synthesized. The reporter strain CDC1551 (*hspX’::GFP*) was treated with across doses of each analog from 200 μM to 0.328 μM. The EC_50_ values of fluorescence inhibition were calculated for each analog to determine their potency. The other chemical properties of the analogs are also included. Kinetic solubility and mouse microsomal stability were both experimentally determined.

| ID# | Compound<br>R = | EC<sub>50</sub><br>(μM) | MW<br>(g/mol) | TPSA<br>(ang<sup>2</sup>) | Ligand<br>Efficiency<br>(LE) | m.p.<br>(°C) | cLogP | Solubility<br>(μM) | Microsomes<br>(% remaining<br>@ 30 minutes) |
| --- | --- | --- | --- | --- | --- | --- | --- | --- | --- |
| HC106F |  | 1.33 | 237.6 | 63 | 0.50 | 183 | 2.2 | >200 | 115% |
| HC106A |  | 2.48 | 272.1 | 63 | 0.41 | 164 | 2.7 | 110 | - |
| MSU-33189 |  | 0.63 | 203.1 | 63 | 0.56 | 183-184 | 1.6 | - | - |
| MSU-39447 |  | 0.61 | 233.2 | 72 | 0.49 | 166-167 | 1.5 | >100 | - |
| MSU-39448 |  | 1.42 | 238.6 | 75 | 0.49 | 158-159 | 1.3 | - | - |
| MSU-39450 |  | 1.95 | 238.6 | 75 | 0.38 | 122-123 | 1.3 | - | - |
| MSU-39446 |  | 0.54 | 221.2 | 63 | 0.52 | 176-177 | 1.7 | >100 | 70% |
| MSU-41464 |  | 1.2 | 282.1 | 67 | 0.51 | 183-184 | 2.3 | - | 109% |
| MSU-39445 |  | 0.75 | 237.6 | 63 | 0.52 | 159 (dec) | 2.1 | >100 | - |
| MSU-39452 |  | 2.47 | 272.1 | 63 | 0.45 | 164 | 2.8 | - | - |
| MSU-41442 |  | 2.08 | 259.3 | 63 | 0.41 | 163 | 3.1 | >100 | - |
| MSU-41443 |  | 11.2 | 279.3 | 63 | 0.32 | 193 | 3.2 | 14 | - |
| MSU-41463 |  | 1.67 | 271.2 | 67 | 0.42 | 178 | 2.4 | - | 84 |
| MSU-41465 |  | 1.16 | 261.2 | 94 | 0.43 | 194 | 1.5 | - | 73 |
| MSU-41462 |  | 1.12 | 217.2 | 67 | 0.45 | 128-130 | 1.3 | - | - |
| MSU-41542 |  | 1.34 | 183.2 | 67 | 0.62 | 187 | 0.9 | - | - |
| MSU-41545 |  | 4.75 | 309.3 | 76 | 0.32 | 164 | 2.9 | - | - |
| MSU-41546 |  | 2.15 | 195.2 | 67 | 0.56 | 182 | 1.1 | - | - |
| MSU-42002 |  | 1.21 | 209.2 | 67 | 0.54 | 180 | 1.4 | - | - |
| MSU-42003 |  | 1.94 | 282.1 | 67 | 0.49 | 157 | 2.3 | - | - |
| MSU-42004 |  | 2.36 | 231.3 | 67 | 0.45 | 165 | 1.7 | - | - |

DosRST is a two-component regulatory system required for Mtb environmental sensing, adaptation, and persistence. By using a fluorescent reporter strain CDC1551 (*hspX’::GFP*), we have previously discovered small molecule inhibitors of DosRST, named HC101A-HC106A, from the whole-cell phenotypic HTS. Here, we report the characterizations of HC104A and HC106A as DosRST inhibitors. Both compounds downregulated genes in the DosR regulon, and inhibited TAG biosynthesis. HC106A also reduced Mtb survival during NRP.

In a UV-visible spectroscopy assay, we observed that HC106A interacts with the heme of sensor kinase DosS. The UV-visible spectrum of HC106A-treated DosS is similar to those of CO-or NO-treated DosS. The overlap between the CO and HC106A spectra supports that HC106A may also directly bind to the heme of DosS. Interestingly, CO activates sensor kinases, whereas HC106A inhibits them. This could be due to the difference in conformational changes induced by CO and HC106A, or binding of HC106A may lock the sensor kinases into an inactive state. Furthermore, the DosS G117L substitution in recombinant DosS blocks the heme exposing channel and provides resistance to HC106A. This means that, similar to artemisinin, this channel is also important for the activity of HC106A. These findings provide additional evidence that the heme-exposing channel in DosS/T can be exploited by small molecules to inhibit the heme from sensing signals and to disrupt signal transduction of a two-component regulatory system. To our knowledge, these two distinct mechanisms of actions are novel mechanisms to inhibit the DosRST pathway.

Mechanistic studies via EMSA indicates that HC104A may function by targeting DosR and inhibiting DosR DNA binding. Virstatin had no effect on DosR DNA binding and no impact on the DosRST signaling in whole cells, showing that although both compounds share a similar structure, they function by distinct mechanisms. From the catalog SAR study, we found that the bromine group is required for the compound. Virstatin does not have the bromine, and the R-group is butyric acid instead of dimethylamine. These two differences are enough to differentiate the activity of the compounds. Moreover, transcriptional profiling shows that the most repressed genes by HC104A are from the DosR regulon, providing additional evidence that HC104A is somewhat selective for specific DosR regulated genes. Interestingly, the genes most downregulated by HC104A, including *hspX, Rv2030c, pfkB*, and *Rv2028c*, are from the same operon under control of *hspX* promoter, which is strongly induced by DosR in hypoxia. This result suggests that HC104A is more specific to target *hspX* operon genes as compared to other DosR regulated genes. This finding leads to the speculation that HC104A may be more efficient to prevent DosR binding to the *hspX* promoter than the other DosR promoters. HC104A may fit better in the pocket in the interface of DosR-*hspX’* complex. This postulation also supports the idea that HC104A may not have an impact on DosR protein dimerization, which would lead to universal downregulation of DosR-regulated genes. Detailed characterizations of HC104A on inhibiting DosR dimerization and promoter binding specificity is the subject of ongoing characterizations.

This study together with the previous report reveals multiple distinct inhibitory mechanisms for the five compounds, HC101A-104A, HC106A. Four out five compounds, HC101A-HC103A and HC106A, are proposed to function by targeting the sensor kinases DosRST. Furthermore, these four compounds are effective at decreasing Mtb survival during NRP in hypoxic-shift down assay. In contrast, the compound targeting DosR, HC104A, had no impact on Mtb survival during NRP. This result is consistent with a previous report that showed only the Δ*dosS* mutant exhibits a survival defect in the C3HeB/FeJ mouse model (22). Notably, the *dosR* mutant strain used in our study is also lacking *dosS* expression due to polar impacts of the deletion, supporting the hypothesis that the reduced survival during NRP may be dependent on DosS. Alternatively, we noted that HC104A only inhibits a portion of the DosR regulon with only 26 genes being downregulated. This finding suggests that the DosR regulated genes that are inhibited by HC104A (Fig. 2, and Supplementary Dataset 1, 3) are not required for DosRST-dependent persistence during NRP or not sufficiently inhibited to have an impact on survival.

The transcriptional profiling data showed off-target impacts in the treated *dosR* mutant strain. These effects may be due to inhibition of DosS/T signaling that functions independent of the response regulator. Several recent studies have demonstrated the occurrence of cross-interactions between histidine kinases and response regulators in Mtb. For instance, Lee *et al.* shows that DosT can interact with the other non-cognate response regulators, including NarL and PrrA (38). Our transcriptional profiling of the compound treated *dosR* mutant strain suggests that some genes downregulated by HC101A, HC103A and HC106A may be DosS-dependent but DosR-independent (Supplementary Fig 9). In the prior study, six genes are similarly regulated between artemisinin and HC103A in the treated *dosR* mutant strain, including Rv0260c that encodes a putative response regulator (27). HC106A and HC103A share four differentially regulated genes in the *dosR* mutant, including *argC, argJ, argB*, and *argF*, which are genes involved in arginine biosynthesis. Our data presented here and in the literature point to the possibility that DosST modulate gene expression independently of DosR.

The synergy studies show significant synergistic interactions between artemisinin, HC102A, HC103A, HC104, or HC106A. Moreover, artemisinin exhibited the greatest synergistic activities with HC102A, HC103A or HC106A, indicating that inhibition of histidine kinases by a second inhibitor can lead to synergistic inhibition of the DosRST pathway. This interaction could be due to both sensor kinases being required for full induction of the DosR regulon, where DosT responds early during hypoxia and DosS further induces the regulon at later times (39), and it is possible that the inhibitors have different affinities for DosS or DosT. Thus, multiple inhibitors could inhibit both DosS and DosT better than an inhibitor alone. Interestingly, artemisinin shows the greatest synergism with HC106A. Both compounds are proposed to target the heme of DosS/T, but through different mechanisms. This finding shows that both inhibitors can enter the channel of DosST to interact with the heme and do so without antagonizing interactions.

The identification of new antibacterial agents and tuberculosis drugs has been associated with the realization that these compounds can occupy a different region of chemical space relative to drugs in most other therapeutic areas. Series HC106 is easily in the range of the characteristics for compounds currently in use or in development as anti-TB compounds. Series HC104 needs further exploration to fully assess its suitability. The physicochemical properties of a drug, such as solubility and permeability, impact its oral bioavailability as these factors influence absorption, distribution, metabolism, and excretion. Series HC106 demonstrates excellent aqueous solubility (Table 2, with compounds generally having solubility >100 µM). The melting points of most of the compounds in the series and the corresponding cLogP are also supportive appropriate solubility. The ability of an anti-TB drug to reach its target site is greatly hampered by the highly impermeable Mtb cell envelope. Both series (HC104 and HC106) demonstrate low micromolar and nanomolar whole cell EC_50_ values, suggesting adequate cell wall permeability. SAR studies of HC106 show that there remain significant opportunities for optimization of HC106 for potency and drug properties and future studies will be focused on such optimizations.

## Methods

### Bacterial strains and growth conditions

Mtb CDC1551, CDC1551 (Δ*dosR*) strains were used in this study. All cultures were grown at 37°C and 5% CO_2_ in 7H9 Middlebrook medium supplemented with 10% OADC (oleic acid albumin dextrose catalase) and 0.05% Tween-80 in standing, vented tissue culture flasks, unless stated otherwise.

***EC*_*50*_ *assays.*** The assay was performed as previously described (27). Briefly, the (*hspX’::GFP*) reporter strain culture was diluted to an OD_600_ of 0.05 in fresh 7H9 media, pH 7.0, and 200 μL of diluted culture was aliquoted in clear-bottom, black, 96-well plates (Corning). Cells were treated with an 8-point (2.5-fold) dilution series ranging from 200 μM – 0.32 μM. For the structure relationship studies for the HC106 series, a 12-point (2.5-fold) dilution series of HC106 analogs ranging from 200 μM – 8.4 nM were used. GFP fluorescence and optical density were measured following 6 d incubation. Percentage fluorescence and growth inhibitions were normalized to a rifampin-positive control (100% inhibition) and DMSO-negative control (0% inhibition). EC_50_ values were calculated for each compound using GraphPad Prism software package (version 6). Each experiment was performed with two technical replicates per plate and two biological replicates, and the error bar represents the s.d. of the biological replicates. Experiments were performed twice with similar results.

### Transcriptional profiling and data analysis

Transcriptional profiling studies were conducted as previously described in Zheng *et al.* (27). Briefly, CDC1551 or CDC1551 (Δ*dosR*) cultures were treated with 40 μM HC104A, HC106A or DMSO control for 6 d. The starting OD_600_ was 0.1 in 8 mL of 7H9 medium in standing T25 vented tissue culture flasks. Bacterial growth consumes oxygen and stimulates the DosRST pathway. The total bacterial RNA from two biological replicates was isolated and prepared for sequencing as previously described (40). The RNA-seq data were processed and analyzed using the SPARTA software package (41). Sequencing data are available at the GEO Database (Accession GSE115892).

### Real time-PCR assays

The vitamin C and NO assays were performed as previously described (27). Briefly, cultures at an OD_600_ of 0.6 were pretreated with 80 μM HC104A, HC106A or a DMSO control for 24 h, and induced with 50 μM DETA-NONOate or 20 mM vitamin C for 2 h. For the HC106A resistance assays, CDC1551 was transformed with the empty replicating plasmid pVV16 or the plasmid expressing *dosS* from the strong *hsp60* promoter (pVV16-DosS), and treated with 20 μM HC106A for 6 d. Total bacterial RNA was isolated and differential gene expression of DosR-regulated genes, including *hspX* and *tgs1*, was quantified. The experiment was performed in three technical replicates and error bars represent the s.d from the mean. The experiment was repeated twice with similar results. To examine Mtb gene expression in macrophages, murine bone-marrow derived macrophages were isolated as previously described (42) and seeded in T75 vented, tissue culture flasks. Macrophages were infected with CDC1551 with multiplicity of infection ratio of 1:20 as previously described (42). After infection, the flasks were treated with 40 μM HC104A or HC106A or DMSO for 48 h, with three individual flasks for each treatment. Total bacterial RNA was isolated after treatment, and the transcripts of DosR-controlled genes (*hspX* and *tgs1*) were quantified in RT-PCR. The experiment was conducted with three biological replicates. The error bar represents the s.d. of the biological replicates.

### TAG biosynthesis

The lipid labelling and TAG TLCs were performed as previously described (27). Briefly, CDC1551 was cultured at an initial OD_600_ of 0.1 and radiolabeled with 8 μCi of [1,2-^14^C] sodium acetate in T25 vented tissue culture flasks. The cultures were treated with 40 μM HC104A, HC106A or DMSO for 6 d at 37°C. CDC1551 (Δ*dosR*) and *dosRS* complement strains were also examined. Total lipids were extracted and ^14^C incorporation was determined by scintillation counting. 20,000 c.p.m. of total lipids were analyzed by TLC using silica gel 60 aluminum sheets (EMD Millipore). To determine TAG accumulation, the lipids were developed in hexane-diethyl ether-acetic acid (80:20:1; vol/vol/vol) solvent system. The TLC was exposed to a phosphor screen for 3 d, and imaged on a Typhoon imager and TAG was quantified using ImageJ software (43). The experiment was repeated twice with similar results, and the error bar represents the s.d. of two biological replicates.

### NRP survival assays

Survival during NRP was examined using the hypoxic shift down assays as previously described (27, 44). Briefly, CDC1551 cells were treated with 40 μM HC104A, HC106A or DMSO control in a 24-well plate (1 mL/well). CDC1551 (Δ*dosR*) and *dosRS* complement strains were also examined. Plates were incubated in an anaerobic chamber (BD GasPak) for 12 d. It took 48 h for cultures to become anaerobic, as monitored by a methylene blue control. Bacterial CFUs were numerated on 7H10 agar plates following incubation. The experiment was repeated twice with similar results.

### DosR protein purification

DosR full length protein was purified as previously described (45). Briefly, the *dosR* gene (Rv3133c) was cloned into pET15b (Novagen Darmstadt, Germany) using the primer set: forward primer 5’-TTT<u>CATATG</u>GTGGTAAAGGTCTTCTTGGTCGATGAC-3’; reverse primer 5’-TTT<u>GGATCC</u>TCATGGTCCATCACCGGGTGG-3’. The His_6_-DosR protein was expressed in *E. coli* BL21(DE3) strain. The culture was grown to OD_600_ 0.5-06, and induced with 1 mM IPTG for 6.5 h at 29°C. The cell pellet was suspended in lysis buffer (20 mM Tris-HCl, pH 8.0, 10% glycerol, 500 mM NaCl, 0.5 mg/ml lysozyme and 0.1 mg/ml PMSF), and incubated at 37°C for 30 min. The soluble fraction of lysate was collected after centrifugation and applied to a TALON metal affinity Co^2+^ column (Clontech). The column was washed twice with washing buffer (20 mM Tris-HCl, pH 8.0, 10% glycerol, 500 mM NaCl) without imidazole, then with 20 mM imidazole. The protein was eluted with the same buffer containing 300 mM imidazole. The fractions with the most DosR protein (as determined by SDS-PAGE) were pooled together for dialysis in 25 mM Tris-HCl, pH 8.0. The final protein concentration was determined using a Qubit kit (Invitrogen).

### Electrophoretic mobility assay

The assay is fluorescence-based using 6-carboxyfluorescein (6FAM) labeled 385 bp probe from the *hspX* promoter. In designing the primer set, 6FAM was added to the 5’ ends of forward and reverse primers. The *hspX* probe was synthesized via PCR using the primer set: forward primer 5’-6FAM-CAACTGCACCGCGCTCTTGATG-3’; reverse primer 5’-6FAM-CATCTCGTCTTCCAGCCGCATCAAC-3’. The probe was purified by Qiagen PCR purification kit. The DosR protein was pre-phosphorylated in 10 μL of phosphorylation buffer (40 mM Tris-HCl, pH 8.0, 5 mM MgCl_2_, 50 mM lithium potassium acetyl phosphate), and incubated at room temperature for 30 min. The protein was then transferred to binding buffer in a final volume of 20 μL (final concentration, 25 mM Tris-HCl, pH 8.0, 0.5 mM EDTA, 20 mM KCl, 6 mM MgCl_2_, 10 nM probe, 1 μg poly-dI-dC (Sigma Aldrich)), and treated with HC104A or an equal volume of DMSO or virstatin (Santa Cruz Biotech). Two different assays were performed. Firstly, different DosR protein concentrations from 0.5 μM to 4 μM were treated with 40 μM. Second, dose response assays were performed with 2 μM DosR treated with different concentrations of HC104A or virstatin from 1 μM to 80 μM. After incubating on ice for 30 min, the reactions were terminated by adding 1 μL 80% glycerol, and loaded on a native 5% Tris/Borate/EDTA (TBE) polyacrylamide gel. The gel was run at 50 V, for 5-6 h at 4°C in 1X TBE buffer, and was imaged using a Typhoon scanner with appropriate filters that can detect florescence at excitation = 495 nm, emission = 520 nm. Binding of the unbound probe was quantified using ImageJ (43). The assay was repeated at least twice with similar results. The error bar represents the s.d. of two biological replicates.

### UV-visible spectroscopy assay

DosS and mutant proteins were purified and analyzed as previously described (27). Briefly, 7.5 μM of recombinant DosS protein was deoxygenated with argon gas in a sealed cuvette. The protein was reduced with 400 μM DTN for 20 min. The reaction was then treated with 100 μM HC106A, 400 μM HC104A, 100 μM CORM-2 (tricarbonyldichlororuthenium (II) dimer) or equal volume of DMSO. The UV-visible spectra were recorded for kinetic changes over 2 h. The experiment was repeated at least twice with similar results.

### Checkerboard synergy studies

The reporter strain CDC1551 (*hspX::GFP*) was treated with pairs of DosRST inhibitors from 50 μM – 0.08 μM in 96-well plates, including HC101A-HC104A and HC106A. GFP fluorescence and OD_600_ were measured after 6 d incubation. The percentage of fluorescence inhibition (FI) and growth inhibition were calculated for each drug pair, with limited growth inhibition observed. The FI data was utilized for further analysis of interactions using CompuSyn software (34). The Combination Index (CI) value was calculated for each drug pair according to the Chou-Talalay method, which is based on the Median-Effect equation derived from the Mass-Action Law principle (35, 46). The resulting CI values provide quantitative determination of drug interactions, including synergism (CI < 1), additive effect (C = 1), and antagonism (C > 1).

### Autophosphorylation assay

The DosS autophosphorylation assay was performed as previously described (27). Recombinant DosS protein was treated with 10 μM, 20 μM or 40 μM of HC104A, or HC106A. DMSO and 40 μM HC103A were also included as positive and negative controls, respectively.

### Kinetic solubility assay

The assay was performed with 7-point (2-fold) dilutions from 200 µM-3.125 µM for HC106 analogs. Mebendazole, benxarotene and aspirin were also included as controls. The drug dilutions were added to PBS, pH 7.4, with the final DMSO concentration of 1%, and incubated at 37 °C for 2 h. The absorbance at 620 nm was measured for each drug dilution to estimate of the compound solubility. Three replicates were examined for each dilution.

## Acknowledgements

The MSU RTSF provided technical support for the RNA-seq library preparation and sequencing. This project was supported by start-up funding and a Molecular Discovery Grant from Michigan State University, AgBioResearch, a grant from the NIH-NIAID (R21AI105687) and Grand Challenges Explorations grants from the Bill and Melinda Gates Foundation (OPP1059227 and OPP1119065).

## Author Contributions

H.Z., E.E. and R.B.A conceived the experiments. H.Z. performed the Mtb physiology and biochemistry experiments. B.A. synthesized the analogs. E.E. directed the medicinal chemistry optimizations. H.Z, E.E. and R.B.A wrote the manuscript.

## Disclosures

RBA is the founder and owner of Tarn Biosciences, Inc., a company that is working to develop new TB drugs.

**Supplementary Methods and Figures**

**Inhibition of *Mycobacterium tuberculosis* DosRST two-component regulatory system signaling by targeting response regulator DNA binding and sensor kinase heme**

## Supplemental Methods

### Experimental procedures for urea formation

#### Formation of acyl chloride 2

To a stirred solution of isoxazole acid **1** (1 eq.) in dry tetrahydrofuran (THF, 0.4 M) under N_2_ atmosphere was added oxalyl chloride (1.5 eq.) dropwise over 5-10 min followed by dimethylformamide (DMF) (cat.) and the reaction mixture was continued to stir at room temperature. Upon completion, the reaction mixture was concentrated into a residue in *vacuo* and the residue was dissolved in THF and concentrated again to ensure the removal of excess oxalyl chloride. The crude acyl chloride **2** was used directly in the next step without further purification.

#### Formation of acyl azide and rearrangement into isocyanate 3

The crude acyl chloride **2** was dissolved in THF (0.4 M) and stirred at room temperature under N_2_ atmosphere. Trimethylsilyl (TMS) azide (2 eq.) was added dropwise over 5 min and stirring was continued. Upon completion of the reaction, the mixture was diluted with ethyl acetate (0.4 M) and quenched with H_2_O (0.4 M). The two layers were separated, and the organic layer was dried over anhydrous Na_2_SO_4_ and filtered. The ethyl acetate solvent was swapped into toluene (0.1 M) by the addition of toluene followed by removal of the ethyl acetate in *vacuo*. Care was taken not to concentrate the toluene. The toluene acyl azide solution was heated at reflux conditions under N_2_ atmosphere for 4 h to give the desired isocyanate **3** which was used as a solution in toluene in the next step.

#### Formation of urea 4

The crude isocyanate solution in toluene was mixed with different amines (1.5 eq.) and stirred at room temperature overnight. Isolation of the ureas was done by diluting the reaction mixture with hexanes, stirring for few hours and filtration of the formed precipitate. The solid material was washed with hexanes and dried under high vacuum. The urea products usually do not require further purifications. All products were analyzed by ^1^H NMR and high-resolution mass spectrometry (HRMS).

#### Synthesis of *<u>MSU-41422</u>* (amide) from acyl chloride 2

Acyl chloride **2** (1 eq.) was dissolved in dichloromethane (DCM, 0.2 M) and 4-chloroaniline (1.2 eq.) was added. The reaction mixture was stirred at room temperature overnight. The reaction mixture was concentrated in *vacuo* and the residue was purified by flash column chromatography.

#### Synthesis of *<u>MSU-39449</u>*

A slurry of 4-chlorophenyl acetic acid (0.10 g, 0.60 mmoles) in 2.0 mL of DCM was treated with oxalyl chloride (0.12 mL, 0.9 mmoles) and 1 drop of DMF. The mixture (with gas evolution) gradually became homogeneous and was stirred for 30 min. The mixture was concentrated in *vacuo*, diluted with 5 mL of DCM and again concentrated in vacuo, the process repeated three times. The resulting residue was again dissolved in DCM (2 mL) and treated with 5-aminoisoxazole (0.030 g., 0.36 mmoles), followed by pyridine (0.48 mL, 0.6 mmoles). The mixture was then allowed to stir overnight. The mixture was then quenched with 1.0 N HCl and extracted with DCM. The organic layers were combined, washed with saturated KHCO_3_, dried with Na_2_SO_4_ and concentrated in *vacuo*. Medium pressure liquid chromatography (SiO_2_, 100% DCM to 3% methanol / DCM) to provide a solid (0.023 g).

#### ^1^H NMR and HRMS data

***MSU# 39444***. ^1^H NMR (500 MHz, DMSO-*d*_6_) δ 10.54 (s, 2H), 8.44 (d, *J* = 1.9 Hz, 2H), 6.11 (d, *J*= 2.0 Hz, 2H). HRMS (ESI) m/z calculated for C_7_H_6_N_4_O_3_ [M+H], 195.0513 found 195.0518.

***MSU# 39449***. ^1^H NMR (500 MHz, DMSO-*d*_6_) δ 11.91 (s, 1H), 8.41 (s, 1H), 7.39 (d, *J* = 2.0 Hz, 2H), 7.35 (d, *J* = 2.0 Hz, 2H), 6.19 (s, 1H), 3.73 (s, 2H). HRMS (ESI) m/z calculated for C_11_H_10_ClN_2_O_2_ [M-H], 235.0269; found 235.1989.

***MSU# 39445***. ^1^H NMR (500 MHz, DMSO-*d*_6_) δ 10.41 (s, 1H), 9.14 (s, 1H), 8.40 (d, *J* = 2.0 Hz, 1H), 7.82 – 7.58 (m, 1H), 7.31 (dd, *J* = 4.9, 1.8 Hz, 2H), 7.07 (dt, *J* = 6.4, 2.3 Hz, 1H), 6.06 (d, *J*= 1.9 Hz, 1H). HRMS (ESI) m/z calculated for C_10_H_9_ClN_3_O_2_ [M+H], 238.0378; found 238.0365.

***MSU# 39452***. ^1^H NMR (500 MHz, DMSO-*d*_6_) δ 10.50 (s, 1H), 9.23 (s, 1H), 8.51 – 8.28 (m, 1H),7.86 (d, *J* = 2.5 Hz, 1H), 7.64 – 7.43 (m, 1H), 7.37 (dd, *J* = 8.9, 2.5 Hz, 1H), 6.26 – 5.80 (m, 1H). HRMS (ESI) m/z calculated for C_10_H_8_Cl_2_N_3_O_2_ [M+H], 271.9989; found 271.9969.

***MSU# 39447***. ^1^H NMR (500 MHz, DMSO-d6) δ 8.83 (s, 1H), 8.43 – 8.29 (m, 1H), 7.36 (d, *J* = 8.4 Hz, 2H), 6.86 (d, *J* = 8.6 Hz, 2H), 6.00 (d, *J* = 1.9 Hz, 1H), 3.70 (s, 3H). HRMS (ESI) m/z calculated for C_11_H_12_N_3_O_3_ [M+H], 234.0874; found 234.0863.

***MSU# 39451***. ^1^H NMR (500 MHz, DMSO-d6) δ 10.13 (s, 1H), 8.34 (d, *J* = 1.9 Hz, 1H), 7.56 – 7.41 (m, 2H), 7.41 – 7.22 (m, 2H), 6.04 (d, *J* = 2.0 Hz, 1H), 3.25 (s, 3H). HRMS (ESI) m/z calculated for C_11_H_11_ClN_3_O_2_ [M+H], 252.0535; found 252.0596.

***MSU# 39453***. ^1^H NMR (500 MHz, DMSO-d6) δ 10.49 (s, 1H), 8.41 (d, *J* = 1.9 Hz, 1H), 7.84 (d, *J* = 8.7 Hz, 1H), 7.27 (dt, *J* = 2.3, 1.1 Hz, 1H), 7.18 (dt, *J* = 8.8, 1.6 Hz, 1H), 6.14 (d, *J* = 1.9 Hz, 1H), 4.13 (dd, *J* = 9.1, 8.2 Hz, 2H), 3.17 (t, *J* = 8.6 Hz, 2H). HRMS (ESI) m/z calculated for C_12_H_11_ClN_3_O_2_ [M+H], 264.0535; found 264.0552.

***MSU# 39448***. ^1^H NMR (500 MHz, DMSO-d6) δ 10.55 (s, 1H), 9.25 (s, 1H), 8.64 – 8.24 (m, 2H), 7.98 (d, *J* = 8.6 Hz, 1H), 7.45 (d, *J* = 8.6 Hz, 1H), 6.08 (d, *J* = 18.1 Hz, 1H). HRMS (ESI) m/z calculated for C_9_H_8_ClN_4_O_2_ [M+H], 239.0331; found 239.0364.

***MSU# 39450***. ^1^H NMR (500 MHz, DMSO-d6) δ 11.00 (s, 1H), 9.66 (s, 1H), 8.42 (d, *J* = 1.9 Hz, 1H), 8.34 (d, *J* = 2.6 Hz, 1H), 7.96 – 7.79 (m, 1H), 7.73 (d, *J* = 8.9 Hz, 1H), 6.10 (q, *J* = 2.7, 2.2 Hz, 2H). HRMS (ESI) m/z calculated for C_9_H_8_ClN_4_O_2_ [M+H], 239.0331; found 239.0356.

***MSU# 39446***. ^1^H NMR (500 MHz, DMSO-*d*_6_) δ 10.24 (s, 1H), 8.90 (s, 1H), 8.38 (d, *J* = 1.9 Hz, 1H), 7.54 – 7.37 (m, 2H), 7.26 – 7.02 (m, 2H), 6.03 (d, *J* = 2.0 Hz, 1H). HRMS (ESI) m/z calculated for C_10_H_9_FN_3_O_2_ [M+H], 222.0674; found 222.0675.

***MSU# 41422***. ^1^H NMR (500 MHz, Chloroform-d) δ 8.41 (d, *J* = 1.8 Hz, 1H), 8.26 (s, 1H), 7.71 – 7.52 (m, 2H), 7.44 – 7.33 (m, 2H), 7.06 (d, *J* = 1.8 Hz, 1H). HRMS (ESI) m/z calculated for C_10_H_8_ClN_2_O_2_ [M+H], 223.0269; found 223.0265.

***MSU# 41324***. ^1^H NMR (500 MHz, DMSO-d6) δ 9.91 (s, 1H), 9.50 (s, 1H), 8.31 (d, *J* = 2.6 Hz, 1H), 7.85 (dd, *J* = 9.0, 2.7 Hz, 1H), 7.67 (d, *J* = 8.9 Hz, 1H), 7.39 – 7.30 (m, 2H). HRMS (ESI) m/z calculated for C_12_H_10_Cl_2_N_3_O [M+H], 282.0196; found 282.0182.

***MSU# 41425***. ^1^H NMR (500 MHz, DMSO-*d*_6_) δ 8.83 (s, 1H), 7.53 – 7.39 (m, 2H), 7.39 – 7.16 (m,2H). HRMS (ESI) m/z calculated for C_13_H_11_Cl_2_N_2_O [M+H], 281.0243; found 281.0258.

***MSU# 41443***. ^1^H NMR (500 MHz, DMSO-*d*_6_) δ 10.36 (d, *J* = 28.4 Hz, 1H), 9.13 (d, *J* = 45.7 Hz, 1H), 8.40 (d, *J* = 1.9 Hz, 1H), 7.70 – 7.59 (m, 5H), 7.59 – 7.49 (m, 3H), 7.43 (t, *J* = 7.7 Hz, 3H), 7.32 (t, *J* = 7.4 Hz, 1H), 6.06 (d, *J* = 1.9 Hz, 1H). HRMS (ESI) m/z calculated for C_16_H_14_N_3_O_2_ [M+H], 280.1081; found 280.1083.

***MSU# 41442***. ^1^H NMR (500 MHz, DMSO-d6) δ 10.15 (s, 1H), 8.77 (s, 1H), 8.38 (d, *J* = 1.9 Hz, 1H), 7.44 – 7.27 (m, 4H), 6.03 (d, *J* = 2.0 Hz, 1H), 1.25 (s, 9H). HRMS (ESI) m/z calculated for C_14_H_18_N_3_O_2_ [M+H], 260.1394; found 260.1406.

***MSU# 33189***. ^1^H NMR (500 MHz, DMSO-*d*_6_) δ 10.20 (s, 1H), 8.85 (s, 1H), 8.39 (d, *J* = 1.9 Hz, 1H), 7.54 – 7.37 (m, 2H), 7.35 – 7.21 (m, 2H), 7.02 (t, *J* = 7.4 Hz, 1H), 6.04 (d, *J* = 1.9 Hz, 1H). HRMS (ESI) m/z calculated for C_10_H_10_N_3_O_2_ [M+H], 204.0768; found 204.0777.

***MSU# 33231***. ^1^H NMR (500 MHz, DMSO-d6) δ 10.29 (s, 1H), 9.01 (s, 1H), 8.39 (d, *J* = 1.9 Hz, 1H), 7.56 – 7.41 (m, 2H), 7.41 – 7.27 (m, 2H), 6.05 (d, *J* = 1.9 Hz, 1H). HRMS (ESI) m/z calculated for C_10_H_9_ClN_3_O_2_ [M+H], 238.0378; found 238.0391.

***MSU# 41462***. ^1^H NMR (500 MHz, DMSO-*d*_6_) δ 10.18 (s, 1H), 8.31 (d, *J* = 2.0 Hz, 1H), 7.43 – 7.07 (m, 3H), 6.92 (s, 1H), 5.94 (d, *J* = 1.9 Hz, 1H), 4.30 (d, *J* = 6.0 Hz, 2H). HRMS (ESI) m/z calculated for C_11_H_12_N_3_O_2_ [M+H], 218.0924 found 218.0956.

***MSU# 41463***. ^1^H NMR (500 MHz, DMSO-*d*_6_) δ 10.39 (s, 1H), 9.29 (s, 1H), 8.41 (d, *J* = 1.9 Hz, 1H), 7.67 (d, *J* = 1.0 Hz, 4H), 6.08 (d, *J* = 1.9 Hz, 1H). HRMS (ESI) m/z calculated for C_11_H_9_F_3_N_3_O_2_ [M+H], 272.0642; found 272.0653.

***MSU# 41464***. ^1^H NMR (500 MHz, DMSO-*d*_6_) δ 10.29 (s, 1H), 9.01 (s, 1H), 8.39 (d, *J* = 1.9 Hz, 2H), 7.50 – 7.45 (m, 1H), 7.45 – 7.40 (m, 1H), 6.05 (s, 1H). HRMS (ESI) m/z calculated for C_10_H_9_BrN_3_O_2_ [M+H], 281.9873; found 281.9876.

***MSU# 41465***. ^1^H NMR (500 MHz, DMSO-*d*_6_) δ 10.38 (s, 1H), 9.27 (s, 1H), 8.41 (d, *J* = 2.0 Hz, 1H), 8.00 – 7.79 (m, 2H), 7.66 – 7.48 (m, 2H), 6.08 (d, *J* = 1.9 Hz, 1H), 3.81 (s, 3H). HRMS (ESI) m/z calculated for C_12_H_12_N_3_O_4_ [M+H], 262.0823; found 262.0827.

***MSU# 41545***. ^1^H NMR (500 MHz, DMSO-*d*_6_) δ 10.17 (s, 1H), 8.71 (s, 1H), 8.37 (d, *J* = 1.9 Hz, 1H), 7.57 – 7.14 (m, 5H), 7.11 – 6.69 (m, 2H), 6.02 (d, *J* = 1.9 Hz, 1H), 5.06 (s, 2H).HRMS (ESI) m/z calculated for C_17_H_16_N_3_O_3_ [M+H], 310.1187; found 310.1178.

***MSU# 41546***. ^1^H NMR (500 MHz, DMSO-*d*_6_) δ 9.69 (s, 1H), 8.30 (d, *J* = 1.9 Hz, 1H), 6.42 (d, *J* = 7.2 Hz, 1H), 5.91 (d, *J* = 1.9 Hz, 1H), 3.92 (h, *J* = 6.7 Hz, 1H), 1.83 (dq, *J* = 12.8, 6.6, 6.0 Hz, 3H), 1.73 – 1.46 (m, 5H), 1.46 – 1.13 (m, 3H). HRMS (ESI) m/z calculated for C_9_H_14_N_3_O_2_ [M+H], 196.1081; found 196.1144.

***MSU# 42002***. ^1^H NMR (500 MHz, DMSO-*d*_6_) δ 9.80 (s, 1H), 8.30 (d, *J* = 1.9 Hz, 1H), 6.34 (d, *J* = 7.8 Hz, 1H), 5.90 (d, *J* = 1.9 Hz, 1H), 3.58 – 3.39 (m, 1H), 1.77 (dt, *J* = 11.1, 3.7 Hz, 1H), 1.64 (dt, *J* = 12.9, 4.1 Hz, 1H), 1.52 (dd, *J* = 10.4, 6.3 Hz, 1H), 1.37 – 0.93 (m, 3H). HRMS (ESI) m/z calculated for C_10_H_16_N_3_O_2_ [M+H], 210.1238; found 210.1282.

***MSU# 42004***. ^1^H NMR (500 MHz, DMSO-*d*_6_) δ 10.07 (s, 2H), 8.31 (d, *J* = 2.0 Hz, 1H), 7.37 – 7.06 (m, 5H), 6.40 (s, 1H), 5.93 (d, *J* = 1.8 Hz, 1H), 3.31 (t, *J* = 7.2 Hz, 2H), 2.74 (t, *J* = 7.2 Hz, 2H). HRMS (ESI) m/z calculated for C_12_H_14_N_3_O_2_ [M+H], 232.1081; found 232.1105.

***MSU# 42003***. ^1^H NMR (500 MHz, DMSO-*d*_6_) δ 10.36 (s, 1H), 9.06 (s, 1H), 8.40 (d, *J* = 1.9 Hz, 1H), 7.83 (t, *J* = 2.0 Hz, 1H), 7.46 – 7.03 (m, 3H), 6.07 (d, *J* = 2.0 Hz, 1H). HRMS (ESI) m/z calculated for C_10_H_9_BrN_3_O_2_ [M+H], 281.9873; found 281.9915.

***MSU# 41542***. ^1^H NMR (500 MHz, DMSO-*d*_6_) δ 10.20 – 9.49 (m, 1H), 8.30 (d, *J* = 1.9 Hz, 1H), 6.44 (t, *J* = 5.9 Hz, 1H), 5.91 (d, *J* = 1.9 Hz, 1H), 2.93 (t, *J* = 6.3 Hz, 2H), 1.68 (dh, *J* = 13.3, 6.7 Hz, 1H), 0.85 (d, *J* = 6.7 Hz, 6H). HRMS (ESI) m/z calculated for C_8_H_14_N_3_O_2_ [M+Na], 206.0899; found 206.0926.

**Supplementary Dataset 1. Differential gene expression data of WT Mtb treated with inhibitors**

**Supplementary Dataset 2. Differential gene expression data of the DosR mutant treated with the inhibitors.**

**Supplementary Dataset 3. Complete gene expression tables for transcriptional profiling experiments.**

**Supplemental Table 1 and Figures 1-9**

---

**Supplemental Table 1.**
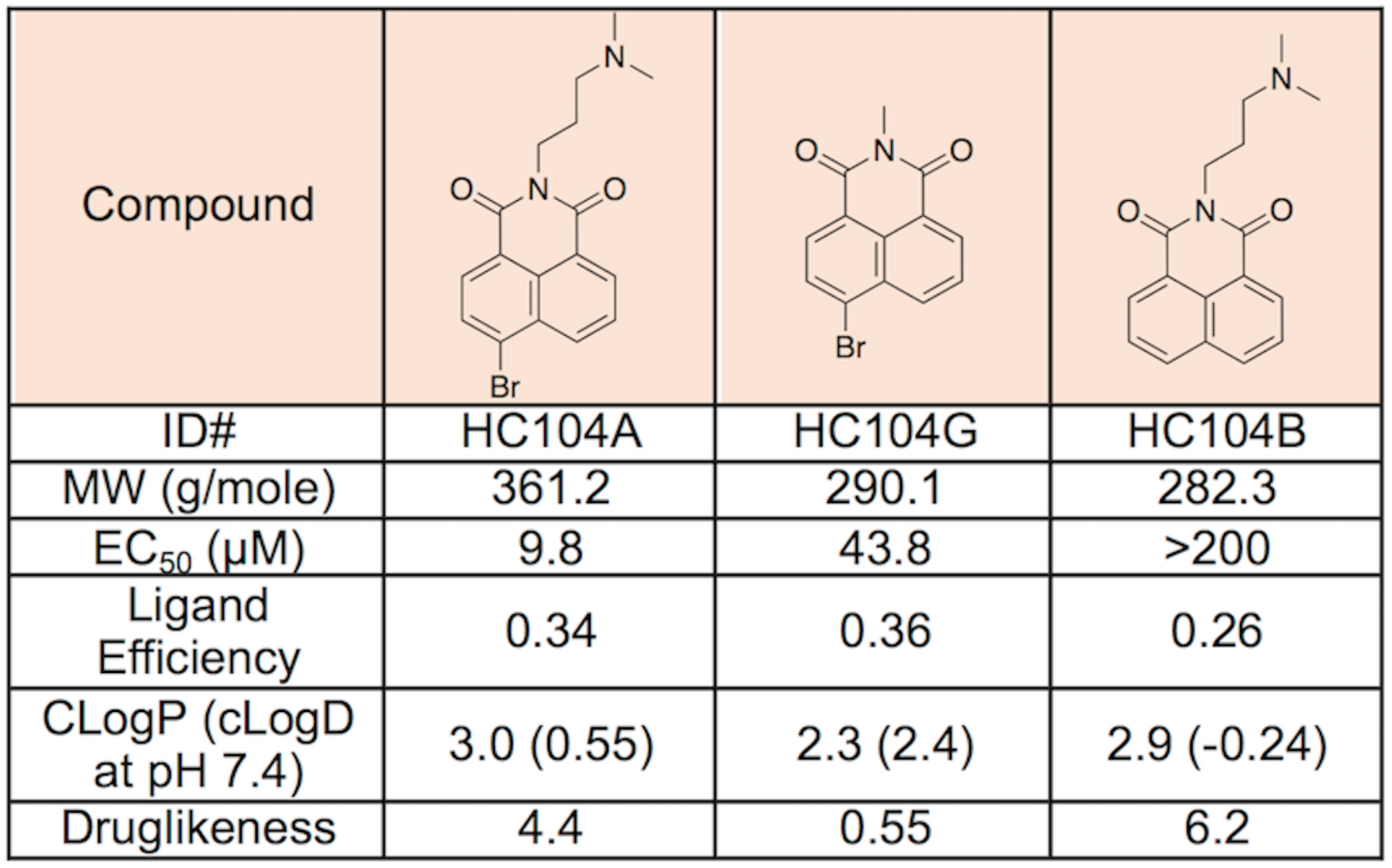
Structure and properties of HC104 series. Catalog structure activity relationship study performed for HC104 analogs with different R-groups. The reporter strain CDC1551 (hspX’::GFP) was treated across doses of each analog from 200 µM to 0.328 µM. The EC50 values of fluorescence inhibition calculated for each analog to determine their potency. The other chemical properties of the analogs are also included.

| Compound |  |  |  |
| --- | --- | --- | --- |
| ID# | HC104A | HC104G | HC104B |
| MW (g/mole) | 361.2 | 290.1 | 282.3 |
| EC<sub>50</sub> (μM) | 9.8 | 43.8 | >200 |
| Ligand Efficiency | 0.34 | 0.36 | 0.26 |
| CLogP (cLogD at pH 7.4) | 3.0 (0.55) | 2.3 (2.4) | 2.9 (-0.24) |
| Druglikeness | 4.4 | 0.55 | 6.2 |

**Supplemental Figure 1.**
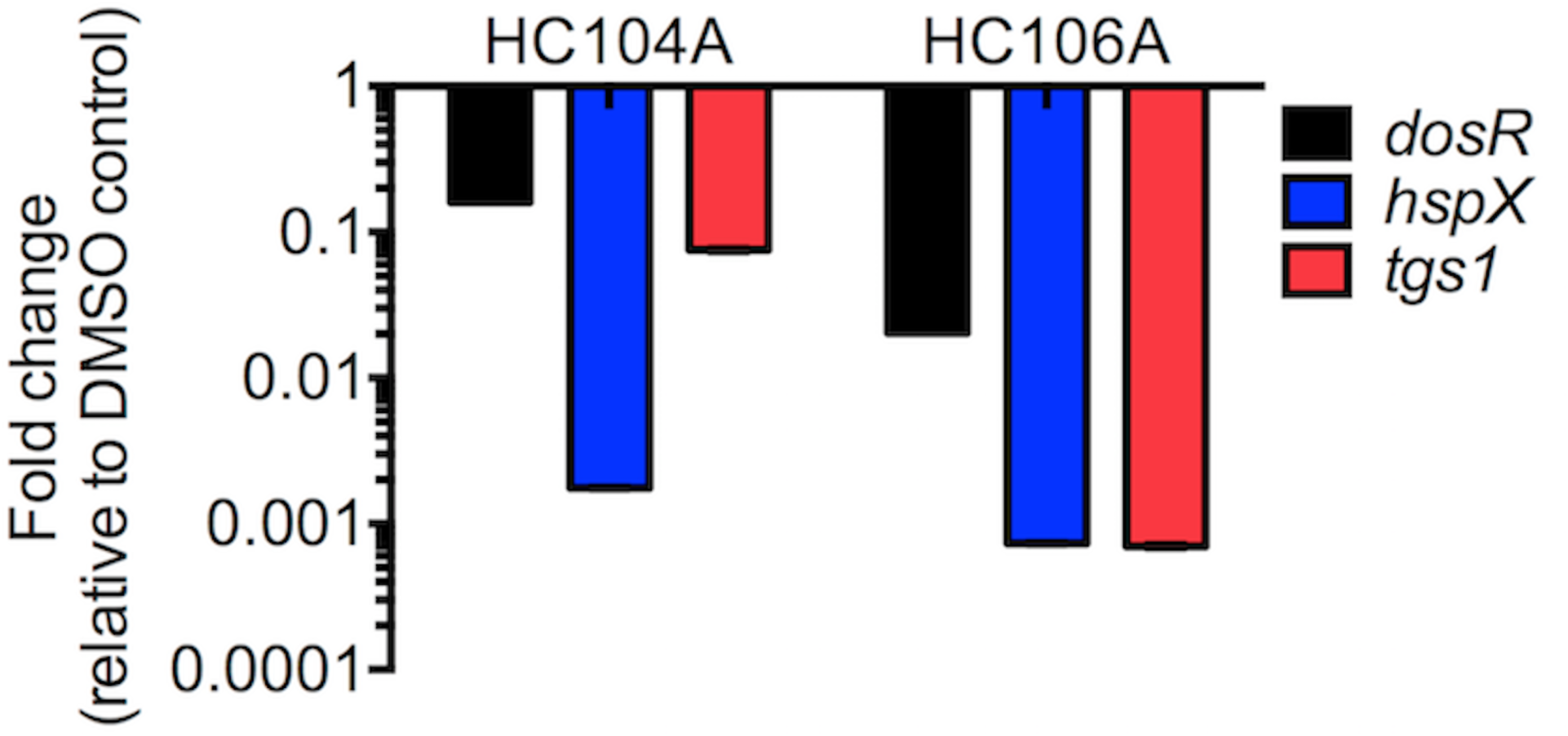
Inhibition of the DosR regulon by HC104 and HC106A during hypoxia. Mtb cells were treated with 40 µM compounds for 6 d, and total bacterial RNA was isolated. The DosR regulated genes, dosR, hspX, and tgs1 were quantified in qRT-PCR. The error bars represent the standard derivation of three replicates. The experiment was repeated at least twice with similar results.

**Supplemental Figure 2.**
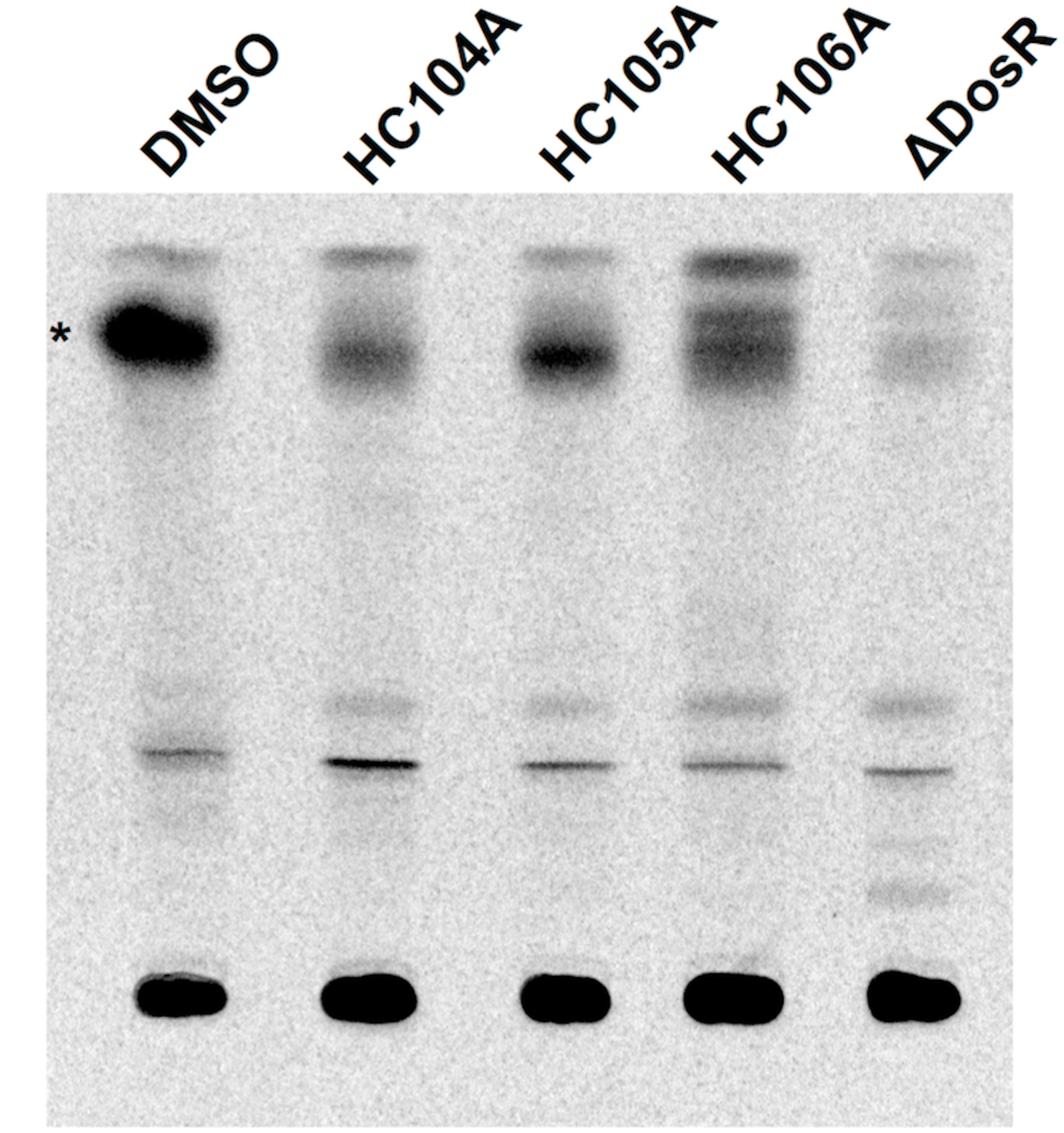
TLC of TAG reduction in Mtb treated with HC104A and HC106A. Mtb cells were treated with 40 µM of the compounds and labeled with [1,2-14C] sodium acetate in T25 vented tissue culture flasks for 6 d. Total lipid was isolated and analyzed BY TLC. The experiment was repeated twice with similar results.

**Supplemental Figure 3.**
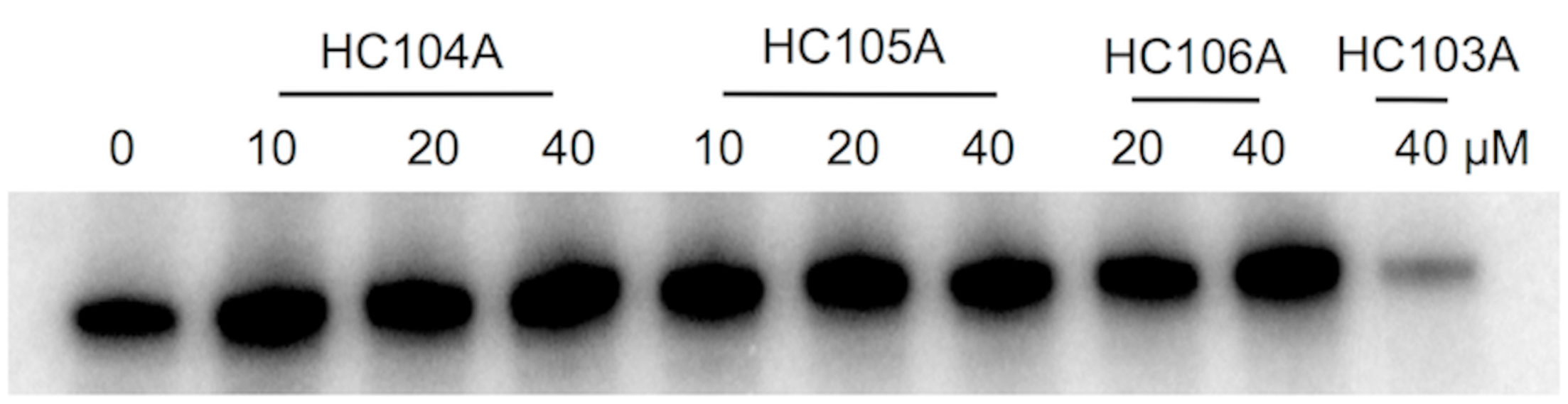
Autoradiograph examining the impact of HC104A and HC106A on DosS autophosphorylation. DosS protein was treated with 10 µM, 20 µM or 40 µM of the compounds, with DMSO and HC103A as positive and negative controls, respectively. The results show that HC104A and HC106A have no effect on DosS autophosphorylation.

**Supplemental Figure 4.**
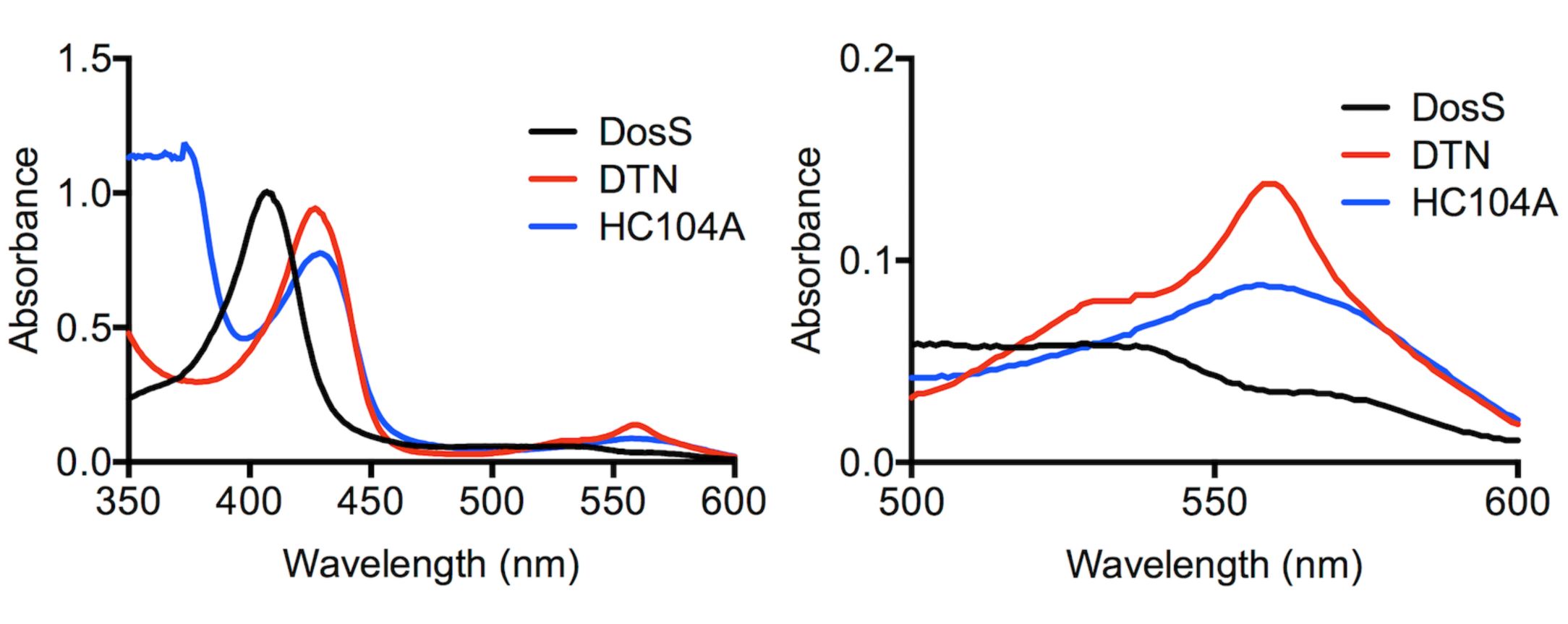
Investigation of interaction between HC104A and DosS. WT DosS treated with 400 µM HC104A shows no impact on shifting of the Soret peak in the UV-visible spectroscopy assay. The experiment was repeated at least twice with similar results.

**Supplemental Figure 5.**
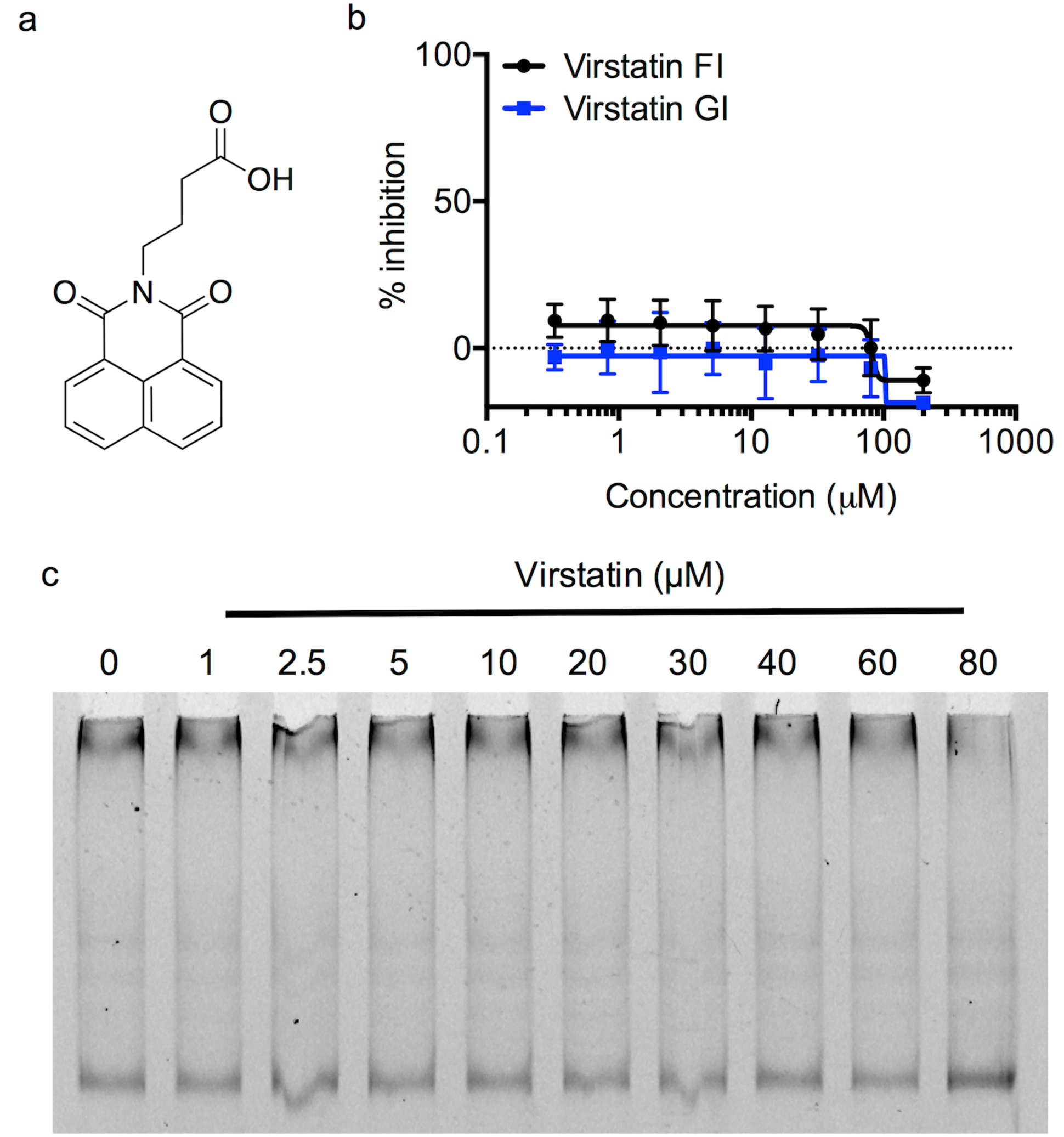
The impact of virstatin on DosR DNA-binding and DosRST signaling in Mtb. (a) Chemical structure of virstatin. (b) Dose-response curve of virstatin shows no effect on inhibition of Mtb DosR-driven GFP fluorescence. (c) DosR protein at 2 µM was treated with 9 point dose response of virstatin from 1 µM to 80 µM. The reactions were analyzed on native PAGE gel. The experiment was repeated at least twice with similar results.

**Supplemental Figure 6.**
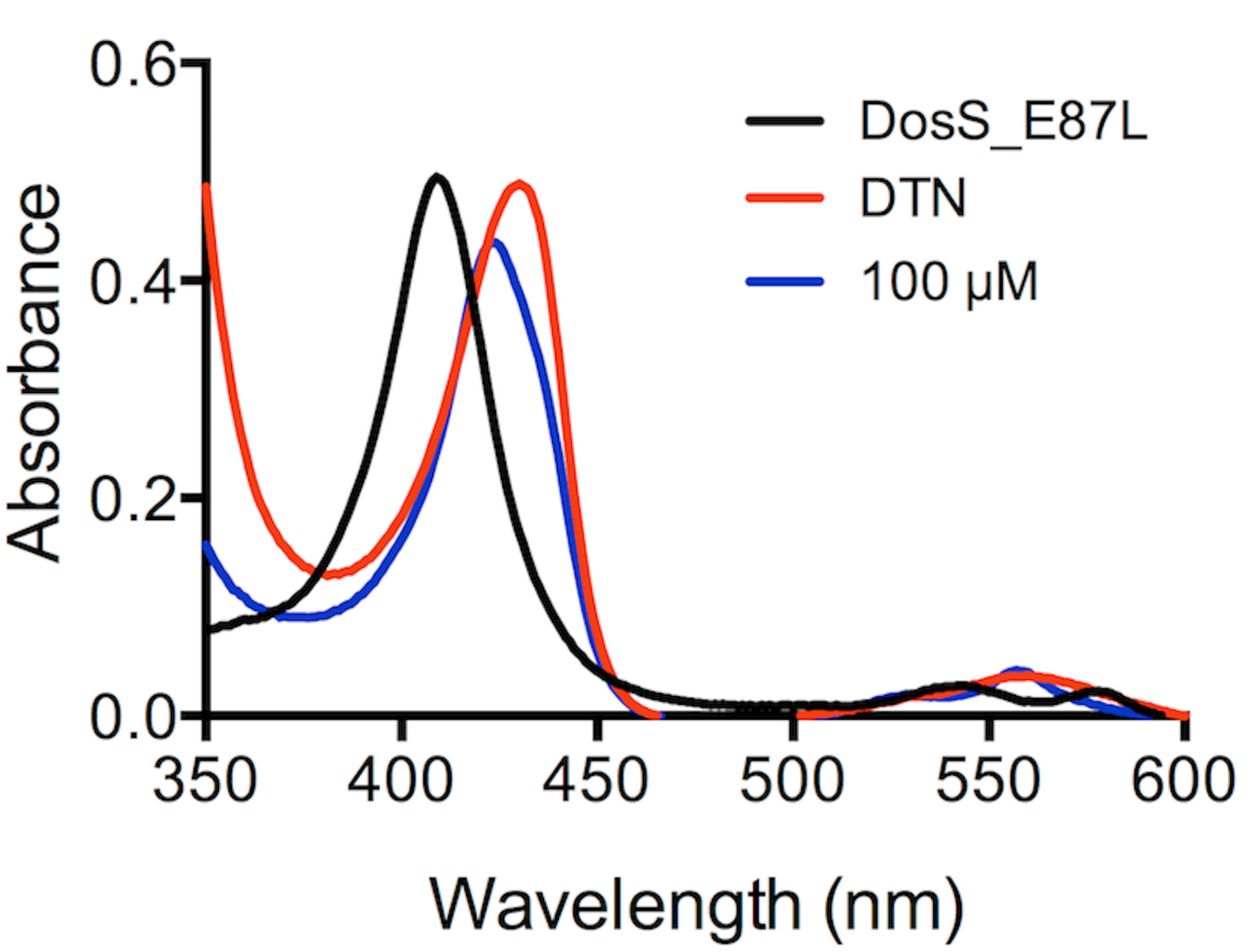
Investigating the interaction between HC106A and DosS heme. DosS E87L protein was treated with 100 µM HC106A after being reduced with DTN. The UV-visible spectra were recorded after each treatment, and showed no change on the overall spectrum compared to WT protein. The experiment was repeated at least twice with similar results.

**Supplemental Figure 7.**
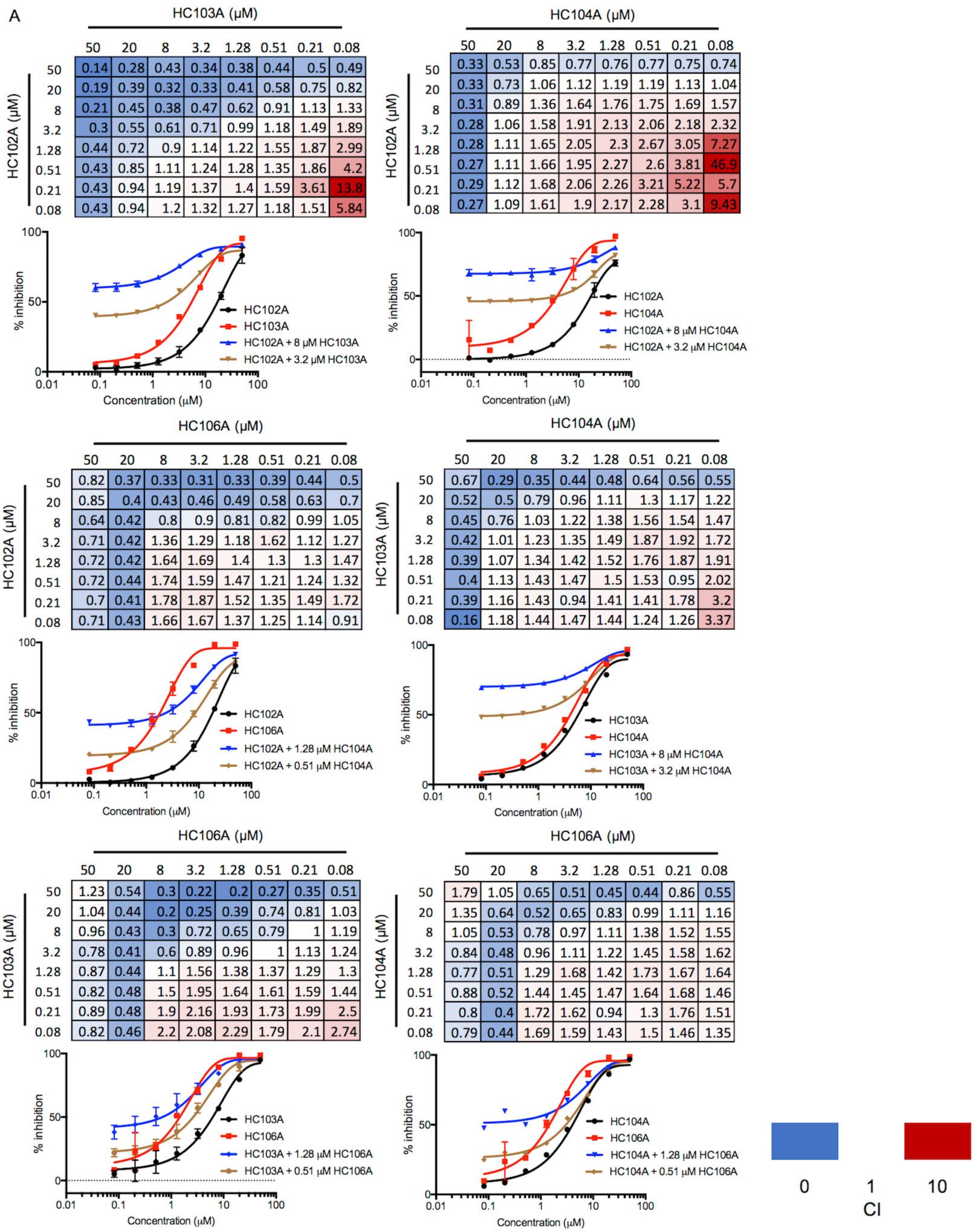
Checkerboard assays examining paired interactions of DosRST inhibitors. CDC1551 (hspX’::GFP) was treated with different combination of two compounds from 50 µM to 0.08 µM. GFP fluorescence measured and used to calculate percentage inhibition. The data were analyzed in the CompuSyn software to determine the combination index (CI) for the panel of each drug combination, including (a) artemisinin and 04A; (b) HC102A and HC103A; (c) HC102A and HC104A; (d) HC102A and HC106A; (e) HC103A and 04A; (f) HC103A and HC106A; (g) HC104A and HC106A. Selected dose response curves are presented to trate synergistic interactions. The experiment was repeated twice with similar results.

**Supplemental Figure 8.**
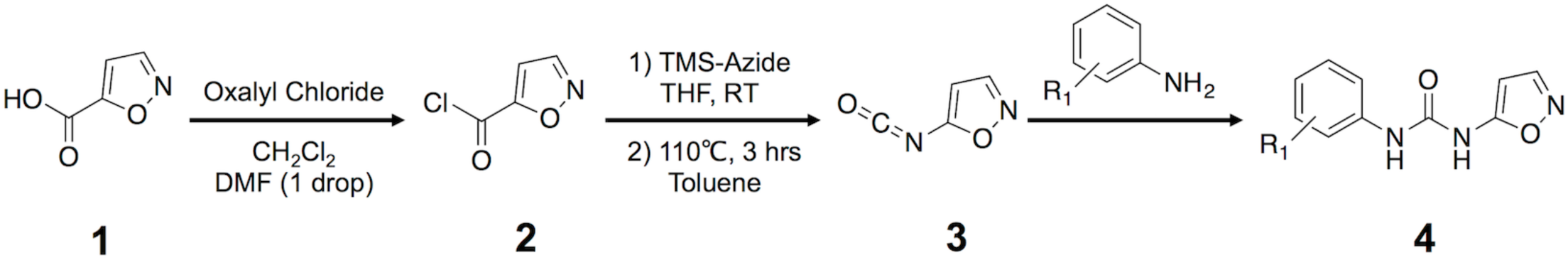
Synthetic scheme for the HC106 analogs.

**Supplemental Figure 9.**
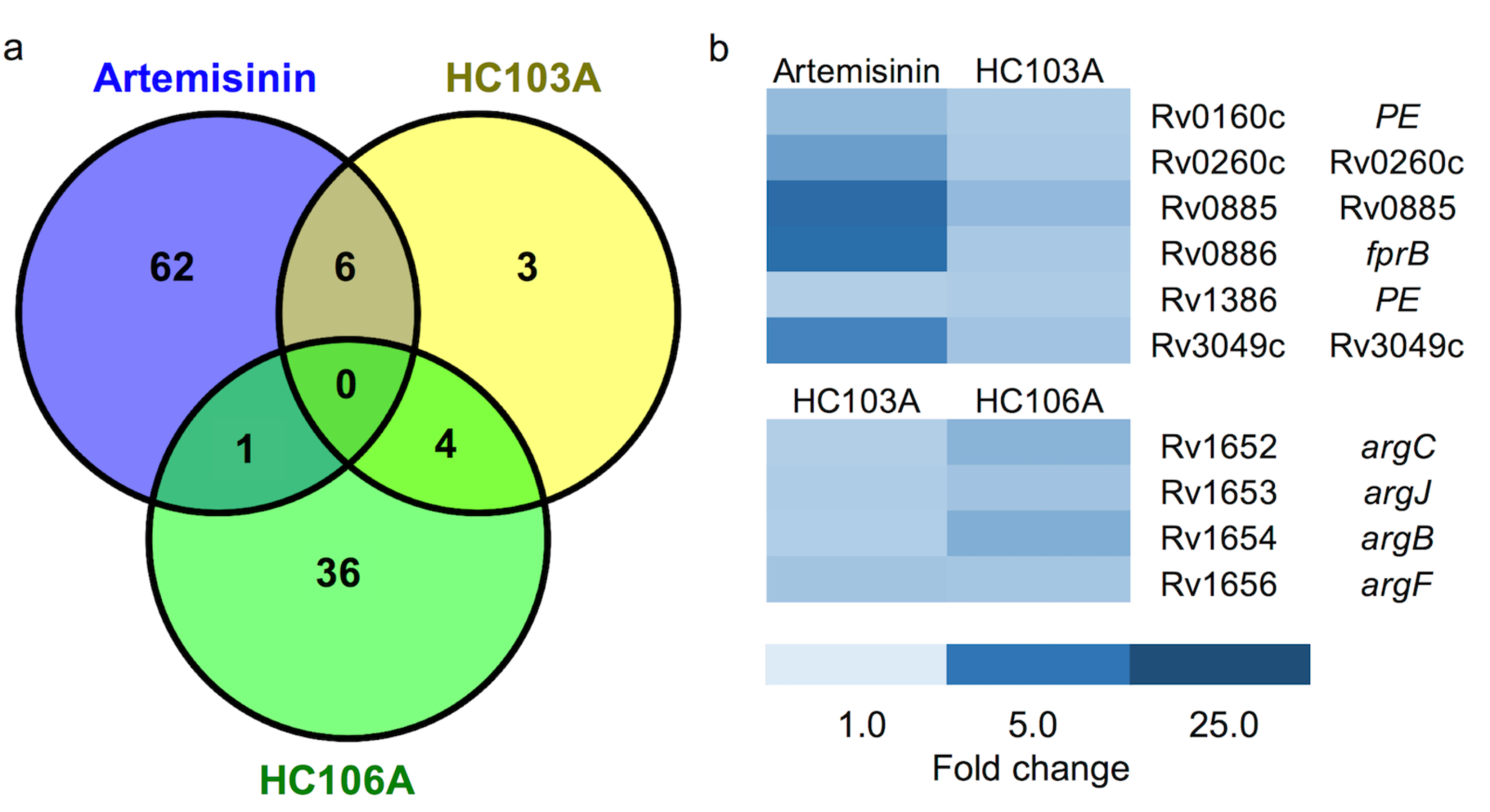
Comparison between artemisinin, HC103 and HC106 for interactions. (a) Venn diagram for the downregulated genes (>2-fold; q< 0.05) of CDC1551 (δdosR) treated with artemisinin, HC103A, or HC106A. (b) Overlap exists between artemisinin and HC103A or HC103A and HC106A differentially expressed genes (downregulated >2X, q<0.05). The heatmaps represent the commonly downregulated genes between the two compounds.

## References

1. Boon C & Dick T (2012) How *Mycobacterium tuberculosis* goes to sleep: the dormancy survival regulator DosR a decade later. Future microbiology 7(4):513–518.

2. Rustad TR, Sherrid AM, Minch KJ, & Sherman DR (2009) Hypoxia: a window into Mycobacterium tuberculosis latency. Cell Microbiol 11(8):1151–1159.

3. Wayne LG & Sohaskey CD (2001) Nonreplicating persistence of mycobacterium tuberculosis. Annu Rev Microbiol 55:139–163.

4. Mehra S, et al. (2015) The DosR Regulon Modulates Adaptive Immunity and Is Essential for Mycobacterium tuberculosis Persistence. American journal of respiratory and critical care medicine 191(10):1185–1196.

5. Hudock TA, et al. (2017) Hypoxia sensing and persistence genes are expressed during the intragranulomatous survival of *Mycobacterium tuberculosis*. American journal of respiratory cell and molecular biology 56(5):637–647.

6. Dasgupta N, et al. (2000) Characterization of a two-component system, devR-devS, of *Mycobacterium tuberculosis*. Tubercle and lung disease : the official journal of the International Union against Tuberculosis and Lung Disease 80(3):141–159.

7. Park HD, et al. (2003) Rv3133c/dosR is a transcription factor that mediates the hypoxic response of Mycobacterium tuberculosis. Mol Microbiol 48(3):833–843.

8. Saini DK, et al. (2004) DevR-DevS is a bona fide two-component system of *Mycobacterium tuberculosis* that is hypoxia-responsive in the absence of the DNA-binding domain of DevR. Microbiology (Reading, England) 150(Pt 4):865–875.

9. Voskuil MI, et al. (2003) Inhibition of respiration by nitric oxide induces a Mycobacterium tuberculosis dormancy program. J Exp Med 198(5):705–713.

10. Kumar A, et al. (2008) Heme oxygenase-1-derived carbon monoxide induces the Mycobacterium tuberculosis dormancy regulon. J Biol Chem 283(26):18032–18039.

11. Lee JM, et al. (2008) O2-and NO-sensing mechanism through the DevSR two-component system in *Mycobacterium smegmatis*. Journal of bacteriology 190(20):6795– 6804.

12. Sousa EH, Tuckerman JR, Gonzalez G, & Gilles-Gonzalez MA (2007) DosT and DevS are oxygen-switched kinases in *Mycobacterium tuberculosis*. Protein science : a publication of the Protein Society 16(8):1708–1719.

13. Kumar A, Toledo JC, Patel RP, Lancaster JR, Jr., & Steyn AJ (2007) Mycobacterium tuberculosis DosS is a redox sensor and DosT is a hypoxia sensor. Proc Natl Acad Sci U S A 104(28):11568–11573.

14. Ioanoviciu A, Yukl ET, Moenne-Loccoz P, & de Montellano PR (2007) DevS, a heme-containing two-component oxygen sensor of *Mycobacterium tuberculosis*. Biochemistry 46(14):4250–4260.

15. Kim MJ, Park KJ, Ko IJ, Kim YM, & Oh JI (2010) Different roles of DosS and DosT in the hypoxic adaptation of Mycobacteria. J Bacteriol 192(19):4868–4875.

16. Rao SP, Alonso S, Rand L, Dick T, & Pethe K (2008) The protonmotive force is required for maintaining ATP homeostasis and viability of hypoxic, nonreplicating Mycobacterium tuberculosis. Proc Natl Acad Sci U S A 105(33):11945–11950.

17. Adams KN, et al. (2011) Drug tolerance in replicating mycobacteria mediated by a macrophage-induced efflux mechanism. Cell 145(1):39–53.

18. Grant SS, Kaufmann BB, Chand NS, Haseley N, & Hung DT (2012) Eradication of bacterial persisters with antibiotic-generated hydroxyl radicals. Proceedings of the National Academy of Sciences of the United States of America 109(30):12147–12152.

19. Leistikow RL, et al. (2010) The Mycobacterium tuberculosis DosR regulon assists in metabolic homeostasis and enables rapid recovery from nonrespiring dormancy. J Bacteriol 192(6):1662–1670.

20. Converse PJ, et al. (2009) Role of the dosR-dosS two-component regulatory system in Mycobacterium tuberculosis virulence in three animal models. Infect Immun 77(3):1230– 1237.

21. Gautam US, Mehra S, & Kaushal D (2015) *in-vivo* gene signatures of *Mycobacterium tuberculosis* in C3HeB/FeJ Mice. PloS one 10(8):e0135208.

22. Gautam US, et al. (2015) DosS is required for the complete virulence of *Mycobacterium tuberculosis* in mice with classical granulomatous lesions. American journal of respiratory cell and molecular biology 52(6):708–716.

23. Deb C, et al. (2009) A novel in vitro multiple-stress dormancy model for Mycobacterium tuberculosis generates a lipid-loaded, drug-tolerant, dormant pathogen. Plos One 4(6):e6077.

24. Basak A, et al. (2017) Antimicrobial peptide-inspired NH125 analogues: bacterial and fungal biofilm-eradicating agents and rapid killers of MRSA persisters. Organic & biomolecular chemistry 15(26):5503–5512.

25. Abramovitch RB (2018) Mycobacterium tuberculosis Reporter Strains as Tools for Drug Discovery and Development. IUBMB Life 70(9):818–825.

26. Johnson BK & Abramovitch RB (2017) Small Molecules That Sabotage Bacterial Virulence. Trends in pharmacological sciences 38(4):339–362.

27. Zheng H, et al. (2017) Inhibitors of Mycobacterium tuberculosis DosRST signaling and persistence. Nature chemical biology 13(2):218–225.

28. Daniel J, et al. (2004) Induction of a novel class of diacylglycerol acyltransferases and triacylglycerol accumulation in *Mycobacterium tuberculosis* as it goes into a dormancy-like state in culture. Journal of bacteriology 186(15):5017–5030.

29. Sirakova TD, et al. (2006) Identification of a diacylglycerol acyltransferase gene involved in accumulation of triacylglycerol in *Mycobacterium tuberculosis* under stress. Microbiology (Reading, England) 152(Pt 9):2717–2725.

30. Baek SH, Li AH, & Sassetti CM (2011) Metabolic regulation of mycobacterial growth and antibiotic sensitivity. PLoS Biol 9(5):e1001065.

31. Hung DT, Shakhnovich EA, Pierson E, & Mekalanos JJ (2005) Small-molecule inhibitor of Vibrio cholerae virulence and intestinal colonization. Science 310(5748):670–674.

32. Podust LM, Ioanoviciu A, & Ortiz de Montellano PR (2008) 2.3 A X-ray structure of the heme-bound GAF domain of sensory histidine kinase DosT of Mycobacterium tuberculosis. Biochemistry 47(47):12523–12531.

33. Cho HY, Cho HJ, Kim YM, Oh JI, & Kang BS (2009) Structural insight into the heme-based redox sensing by DosS from Mycobacterium tuberculosis. J Biol Chem 284(19):13057–13067.

34. Chou T & Martin N (2005) CompuSyn for drug combinations: PC software and user’s guide: a computer program for quantitation of synergism and antagonism in drug combinations, and the determination of IC50 and ED50 and LD50 values).

35. Chou TC & Talalay P (1984) Quantitative analysis of dose-effect relationships: the combined effects of multiple drugs or enzyme inhibitors. Adv Enzyme Regul 22:27–55.

36. Boulton AJ & Katritzky AR (1961) The tautomerism of heteroaromatic compounds with five-membered rings—I: 5-hydroxyisoxazoles-isoxazol-5-ones. Tetrahedron 12(1):41–50.

37. Topliss JG (1972) Utilization of operational schemes for analog synthesis in drug design. Journal of medicinal chemistry 15(10):1006–1011.

38. Lee HN, Jung KE, Ko IJ, Baik HS, & Oh JI (2012) Protein-protein interactions between histidine kinases and response regulators of *Mycobacterium tuberculosis* H37Rv. Journal of microbiology (Seoul, Korea) 50(2):270–277.

39. Honaker RW, Leistikow RL, Bartek IL, & Voskuil MI (2009) Unique roles of DosT and DosS in DosR regulon induction and Mycobacterium tuberculosis dormancy. Infect Immun 77(8):3258–3263.

40. Baker JJ, Johnson BK, & Abramovitch RB (2014) Slow growth of Mycobacterium tuberculosis at acidic pH is regulated by phoPR and host-associated carbon sources. Mol Microbiol 94(1):56–69.

41. Johnson BK, Scholz MB, Teal TK, & Abramovitch RB (2016) SPARTA: Simple Program for Automated reference-based bacterial RNA-seq Transcriptome Analysis. BMC bioinformatics 17:66.

42. Johnson BK & Abramovitch RB (2015) Macrophage Infection Models for Mycobacterium tuberculosis. Methods Mol Biol 1285:329–341.

43. Schneider CA, Rasband WS, & Eliceiri KW (2012) NIH Image to ImageJ: 25 years of image analysis. Nature methods 9(7):671–675.

44. Mak PA, et al. (2012) A high-throughput screen to identify inhibitors of ATP homeostasis in non-replicating Mycobacterium tuberculosis. ACS chemical biology 7(7):1190–1197.

45. Vashist A, Prithvi Raj D, Gupta UD, Bhat R, & Tyagi JS (2016) The alpha10 helix of DevR, the *Mycobacterium tuberculosis* dormancy response regulator, regulates its DNA binding and activity. The FEBS journal 283(7):1286–1299.

46. Chou TC (2010) Drug combination studies and their synergy quantification using the Chou-Talalay method. Cancer research 70(2):440–446.

